# Persistent microbial material contributes to Alzheimer disease and is targetable by vaccination

**DOI:** 10.64898/2026.07.20.739171

**Authors:** Evan C. Marcet, Mariana Vinacur, Tanweer Zaidi, Nhi Nguyen, Anna Augart-Welwood, Grace Su, Dylan Landau, Elijah LaVancher, Artem Arkhangelskiy, Liam H. Power, Marilyn F. Kelly, Casey R. Stein, Eytan Carolan, Barbara Caldarone, Cynthia A. Lemere, Caroline Wasén, Laura M. Cox, Dana Cairns, David L. Kaplan, Gerald B. Pier, Colette Cywes-Bentley

**Affiliations:** Department of Biomedical Engineering, Tufts University; Medford, MA, 02155, USA; Division of Infectious Diseases, Department of Medicine, Brigham and Women’s Hospital, Harvard Medical School; Boston, MA, 02115, USA; Boston College; Chestnut Hill, MA, 02467, USA; Mouse Behavior Core Facility, Harvard Medical School; Boston, MA, 02115, USA; Department of Neurology, Ann Romney Center for Neurologic Diseases at Brigham and Women’s Hospital and Harvard Medical School; Boston, MA, 02115, USA

## Abstract

Chronic neuroinflammation is increasingly recognized as a contributor to Alzheimer disease, yet the upstream stimuli that sustain it remain poorly defined. We investigated whether persistent microbial material contributes to Alzheimer disease using the conserved microbial polysaccharide poly-*N*-acetylglucosamine (PNAG). PNAG-containing microbial material colocalized with amyloid plaques in human Alzheimer disease brain tissue. In fully human neuronal and three-dimensional brain models, purified PNAG and PNAG-containing microbial vesicles activated Toll-like receptor 2-dependent inflammasome signaling and promoted amyloid-β and phosphorylated tau accumulation. Vaccination targeting PNAG improved cognition, reduced glial activation and amyloid pathology, remodeled amyloid processing, and preserved gut microbial community structure in APP/PS1 mice. These findings identify persistent microbial material as an upstream contributor to Alzheimer disease-associated neuroinflammation and a potential therapeutic target.

## Main Text

Alzheimer disease (AD) is characterized by progressive neurodegeneration accompanied by amyloid-β (Aβ) deposition, tau pathology, and chronic neuroinflammation (*1–3*). Although increasing evidence implicates innate immune activation in disease progression, the upstream stimuli that sustain chronic inflammatory signaling remain poorly defined (*4–6*). Microbial products, alterations in the intestinal microbiota, and systemic inflammatory responses have each been associated with AD (*7–9*), but whether conserved microbial material directly contributes to neurodegeneration has remained unresolved.

Persistent microbial material—including extracellular vesicles, cell wall components, polysaccharides, and other microbial products—can disseminate systemically, persist after microbial clearance, and activate innate immune pathways through pattern-recognition receptors (*5–12*). Poly-*N*-acetylglucosamine (PNAG) is a highly conserved microbial polysaccharide expressed by numerous bacterial and fungal species but absent from mammalian cells (*13–19*). Because PNAG is a common constituent of microbial material, including bacterial extracellular vesicles and cell wall fragments (*17,19*), it provides an experimentally tractable target for determining whether microbial material persisting in the brain contributes directly to chronic neuroinflammation.

Here, we investigated whether PNAG-containing microbial material contributes to AD pathology and whether immune targeting of this conserved microbial antigen modifies disease-relevant outcomes. We show that PNAG-containing microbial material colocalizes with amyloid plaques in human AD brain tissue and that purified PNAG and PNAG-containing microbial vesicles activate TLR2-dependent inflammasome signaling while promoting Aβ accumulation and tau phosphorylation in fully human neuronal systems. Vaccination targeting PNAG attenuated glial activation, reduced amyloid pathology, shifted amyloid processing toward less aggregation-prone Aβ isoforms, and preserved cognitive function in APP/PS1 mice without altering gut microbial community composition. These experimental results support the hypothesis that persistent microbial material functions as an upstream driver of AD-associated neuroinflammation and that immune targeting of the conserved microbial antigen PNAG modifies disease progression (Fig.1).

**Figure 1.**
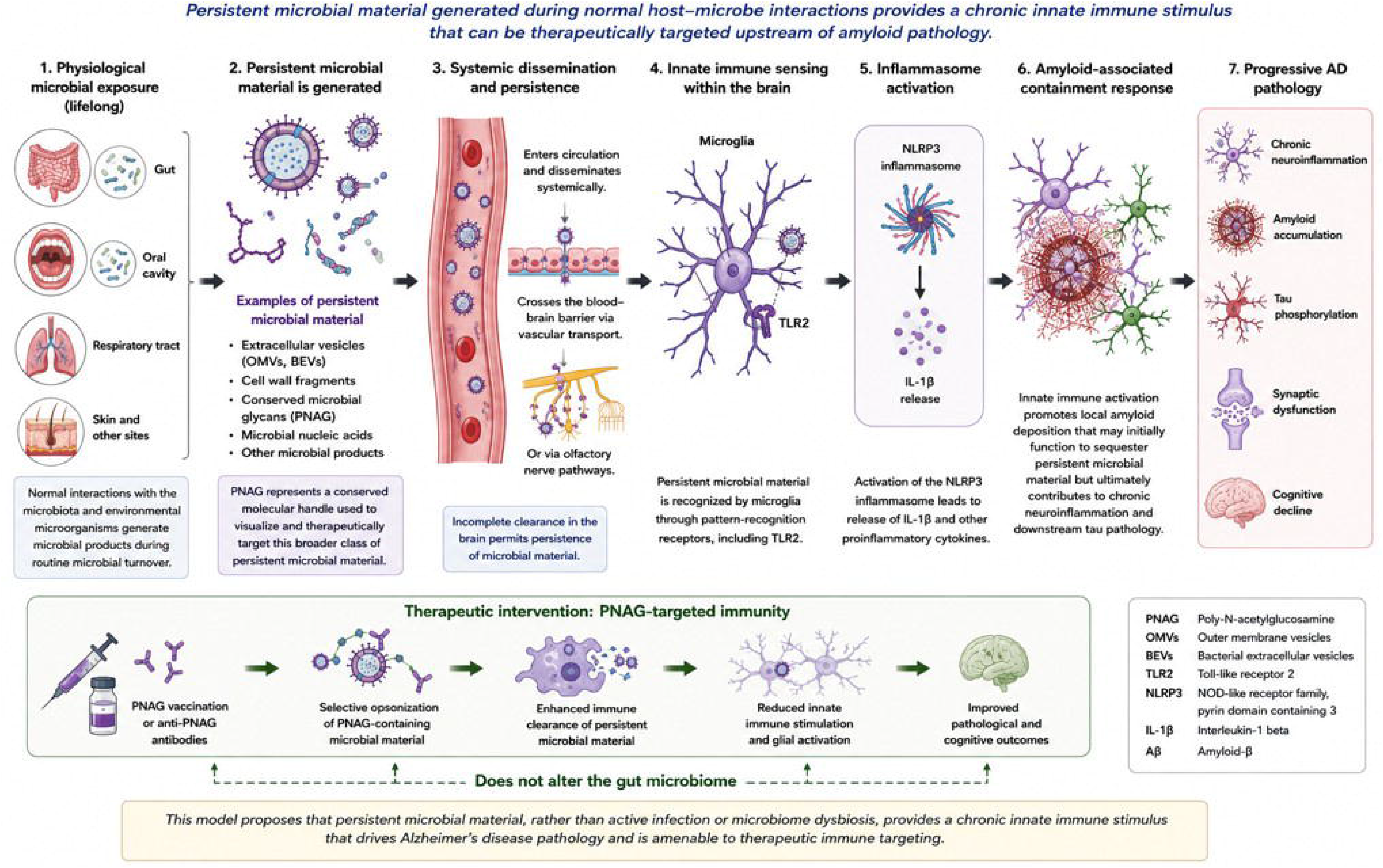
Proposed model: Persistent microbial material as an upstream therapeutic target in Alzheimer’s.

### Persistent PNAG-containing microbial material accumulates in association with amyloid plaques in human AD brain

To determine whether persistent microbial material is present in human AD brain, we examined postmortem tissue from patients with AD and age-matched non-AD controls by confocal immunofluorescence microscopy using the PNAG-specific monoclonal antibody F598 (*16*) together with the Aβ antibody 6E10 (Fig. 2A). PNAG-containing microbial material was abundant throughout AD brain regions commonly affected by disease, including the hippocampus, cortex, and substantia nigra, where it localized within or immediately adjacent to Aβ plaques. In contrast, PNAG immunoreactivity was minimal in non-AD control brains and showed little association with the occasional Aβ deposits present in these tissues (Fig. 2A). Quantitative image analysis demonstrated significantly greater spatial colocalization of PNAG-containing microbial material with Aβ in AD than in control brains (Welch’s *t* test, P < 0.0001; d*_Cohen_*= 6.19; 95% CI, 3.743–8.629; Fig. 2B). To confirm antibody specificity, tissue sections were treated with the PNAG-specific glycosyl hydrolase Dispersin B (*20*), which abolished PNAG immunoreactivity while preserving Aβ staining (fig. S1), confirming that the detected signal represented authentic PNAG-containing microbial material. These findings identify persistent PNAG-containing microbial material as a consistent component of human AD pathology and demonstrate its close spatial association with amyloid plaques.

**Figure 2.**
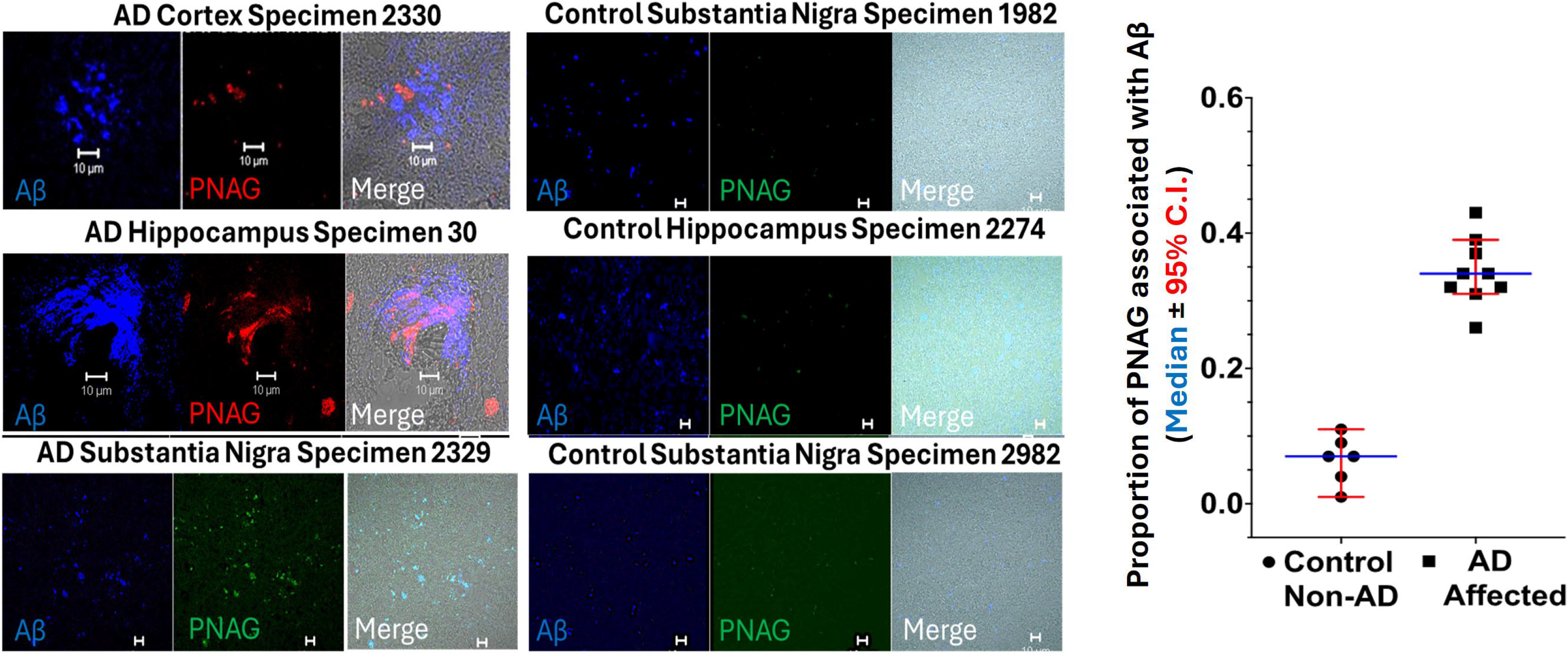
Persistent microbial material colocalizes with amyloid plaques in human Alzheimer disease brain. **(A)** Representative confocal images of postmortem Alzheimer disease (AD) and age-matched non-AD control brain tissue immunostained for amyloid-β (Aβ, blue) and PNAG-containing microbial material (red). PNAG-containing microbial material colocalizes with Aβ plaques in AD brain but is minimal in non-AD controls. Scale bars, 10 μm.**(B)** Quantification of spatial colocalization between PNAG-containing microbial material and Aβ in AD and control brains. Data are median ± 95% CI. Welch’s *t* test, P < 0.0001; d*_Cohen_* = 6.19; 95% CI, 3.743–8.629; large effect size.

### Persistent microbial material directly initiates innate immune activation and AD-associated pathology in human neuronal cultures

To determine whether persistent microbial material is sufficient to initiate AD-associated inflammatory responses, human induced neural stem cell (hiNSC)-derived neuronal cultures (*21,22*) were exposed for four days to purified PNAG isolated from either *Staphylococcus aureus* or *Acinetobacter baumannii* (Fig. 3A) (*13,14*). Purified PNAG from both bacterial species induced robust intracellular accumulation of Aβ that closely colocalized with PNAG, whereas PBS-treated cultures exhibited little detectable Aβ (Fig. 3A).

**Figure 3.**
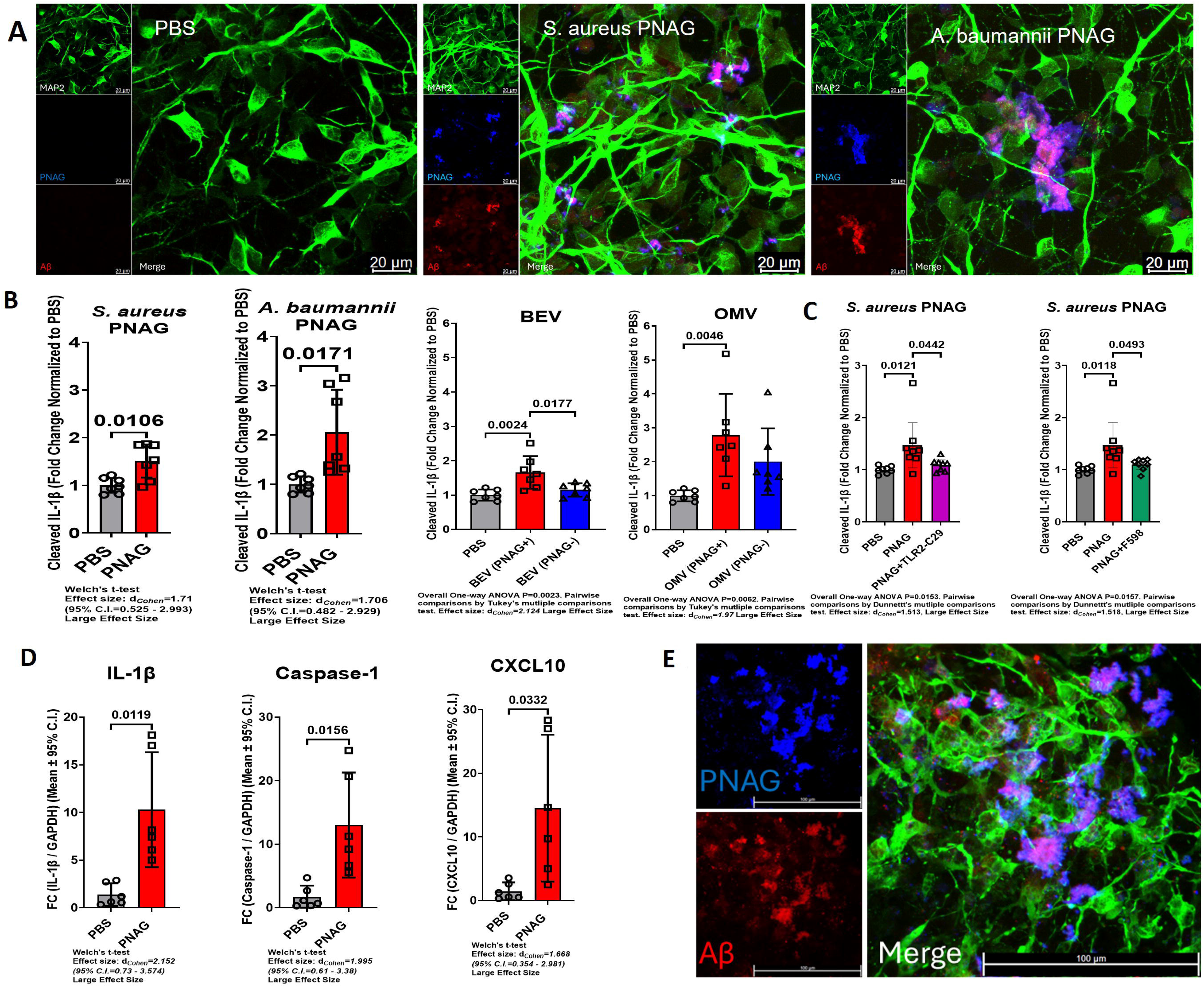
Purified PNAG and PNAG-positive microbial vesicles initiate innate immune activation in fully human neuronal systems. **(A)** Representative confocal images of hiNSC-derived neuronal cultures treated for 4 days with PBS or purified *Staphylococcus aureus* or *Acinetobacter baumannii* PNAG and stained for MAP2 (green), PNAG (blue), and amyloid-β (Aβ, red). Scale bars, 20 μm. **(B)** Bioactive IL-1β released from neuronal cultures following exposure to purified PNAG, PNAG-positive or PNAG-deficient bacterial extracellular vesicles (BEVs), and outer membrane vesicles (OMVs). **(C)** Effects of Toll-like receptor 2 (TLR2) inhibition and the PNAG-specific human monoclonal antibody F598 on PNAG-induced IL-1β production. **(D)** Relative expression of IL1β, CASP1, and CXCL10 following PNAG exposure. **(E)** Representative confocal images of a three-dimensional human neuron–astrocyte–microglia (NAM) culture treated with purified PNAG and stained for PNAG (blue), Aβ (red), and MAP2 (green). Scale bars, 100 μm. Data are mean ± 95% CI.

Purified PNAG also significantly increased secretion of bioactive IL-1β compared with vehicle-treated cultures (*S. aureus*, P = 0.0106; *A. baumannii*, P = 0.0171; Fig. 3B). Similar responses were induced by PNAG-positive bacterial extracellular vesicles (BEVs) and outer membrane vesicles (OMVs), whereas PNAG-deficient vesicles elicited substantially weaker responses (Fig. 3B). Pharmacologic inhibition of Toll-like receptor 2 (TLR2) significantly reduced PNAG-induced IL-1β production (P = 0.0442; Fig. 3C), and the PNAG-specific fully human monoclonal antibody F598 (*14,19,23*) similarly attenuated IL-1β release (P = 0.0493; Fig. 3C), demonstrating that innate immune activation is mediated through TLR2-dependent recognition of PNAG-containing microbial material. Consistent with inflammasome activation, PNAG significantly increased expression of IL1β, CASP1, and CXCL10 (Fig. 3D). Quantitative ELISA confirmed increased Aβ42 production following PNAG exposure (P = 0.0346; Fig. 4C). In complementary experiments, human induced pluripotent stem cell-derived neurons (*24*) exposed to PNAG exhibited increased phosphorylation of tau detected with the AT8 antibody (P = 0.0192; Fig. 4C). Together, these findings demonstrate that persistent microbial material is sufficient to activate TLR2-dependent innate immune signaling while promoting the accumulation of the principal pathological protein hallmarks of AD in fully human neuronal systems.

**Figure 4.**
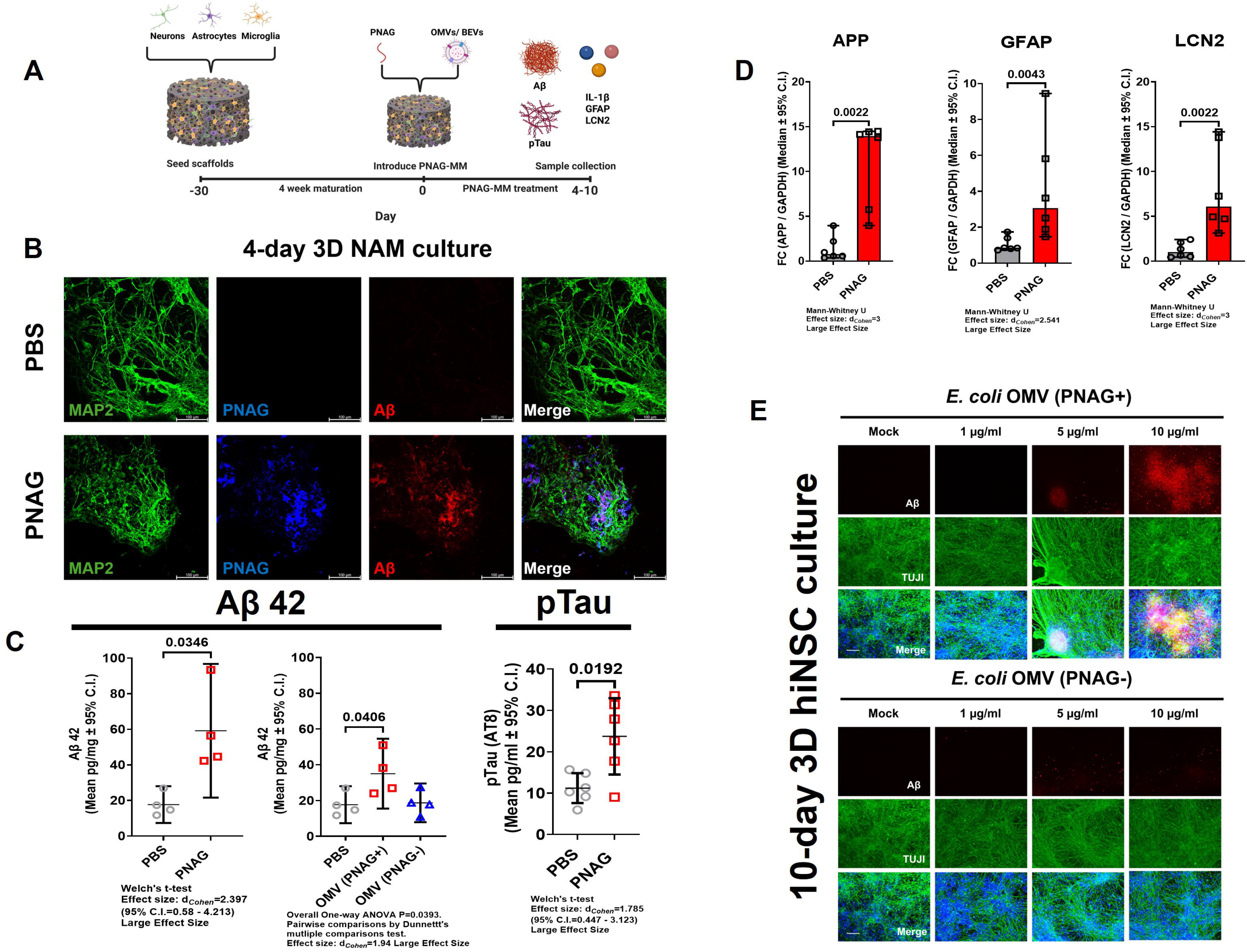
Persistent microbial material reproduces AD-associated pathology in three-dimensional human brain models. **(A)** Experimental design for three-dimensional (3D) human neuron–astrocyte–microglia (NAM) cultures. Silk scaffolds were seeded with human neurons, astrocytes, and microglia, matured for 4 weeks, and exposed to purified PNAG or PNAG-containing bacterial extracellular vesicles (BEVs) or outer membrane vesicles (OMVs). Samples were collected 4–10 days after treatment. **(B)** Representative confocal images of 3D NAM cultures treated with PBS or purified PNAG and stained for MAP2 (green), PNAG (blue), and amyloid-β (Aβ, red). Scale bars, 100 μm. **(C)** Quantification of Aβ42 and phosphorylated tau (pTau; AT8) following exposure to purified PNAG or *Escherichia coli* OMVs. **(D)** Relative expression of APP, GFAP, and LCN2 following PNAG exposure. **(E)** Representative confocal images of 10-day 3D human neuronal cultures exposed to increasing concentrations of PNAG-positive or PNAG-deficient *E. coli* OMVs and stained for TUJ1 (green) and Aβ (red). Scale bars, 100 μm.

### Persistent microbial material reproduces AD-associated pathology in physiologically relevant human brain models

To determine whether the pathological responses induced by purified PNAG were reproduced by physiologically relevant microbial material, differentiated human neuronal cultures and engineered three-dimensional human brain models were exposed to PNAG-positive or PNAG-deficient bacterial outer membrane vesicles (OMVs) (*25*) according to the experimental design shown in Fig. 4A. Exposure to PNAG-positive OMVs significantly increased Aβ42 production compared with vehicle-treated controls (P = 0.0406), whereas PNAG-deficient OMVs induced substantially less Aβ42 accumulation (Fig. 4C). PNAG-positive OMVs also increased expression of APP, GFAP, and LCN2 compared with PNAG-deficient vesicles (Fig. 4D), demonstrating activation of AD-associated molecular pathways. These responses were maintained in physiologically relevant three-dimensional human brain models. Mature neuron–astrocyte–microglia (NAM) cultures (*21*) treated with purified *Staphylococcus aureus* PNAG exhibited robust Aβ accumulation together with spatial colocalization of persistent microbial material and amyloid deposits (Fig. 4B), closely recapitulating the spatial relationship observed in postmortem human AD brain tissue (Fig. 2A). Likewise, collagen-embedded three-dimensional human neuronal cultures (*22*) developed prominent fibrillar Aβ accumulation following exposure to PNAG-positive OMVs, whereas PNAG-deficient OMVs induced substantially less pathology (Fig. 4E). Together, these findings demonstrate that physiologically relevant microbial vesicles reproduce the molecular and pathological responses induced by purified PNAG in complex human brain models, establishing that persistent microbial material remains biologically active in its natural vesicular context.

### Immune targeting of persistent microbial material preserves cognitive function in APP/PS1 mice

APP/PS1 mice were immunized with a PNAG–tetanus toxoid (PNAG-TT) conjugate vaccine previously shown to be safe and immunogenic in Phase 1 clinical studies (Fig. 5A) (*23*). Control animals received tetanus toxoid (TT) alone. PNAG-TT vaccination elicited robust anti-PNAG IgG responses that were maintained throughout the study, whereas TT-immunized mice developed no detectable PNAG-specific antibodies (fig. 5B; fig. S3). Importantly, mice were not challenged with exogenous PNAG or experimental microbial inocula, such that the observed effects reflect immune targeting of naturally encountered persistent microbial material. APP/PS1 mice vaccinated beginning at either 5 weeks or 5 months of age underwent behavioral assessment at 13 or 15 months, respectively. During Water T-maze testing, both early- and delayed-vaccinated mice performed significantly better than TT-vaccinated APP/PS1 controls during acquisition and reversal learning (both P < 0.0001; d*_Cohen_* > 1.25; Fig. 5C). Performance of vaccinated APP/PS1 mice was indistinguishable from non-transgenic littermates (P > 0.1175). In open-field testing, PNAG-vaccinated APP/PS1 mice exhibited reduced hyperlocomotor activity compared with TT-vaccinated APP/PS1 mice, with exploratory behavior comparable to that of non-transgenic controls (Fig. 5D). Together, these findings demonstrate that immune targeting of naturally encountered persistent microbial material preserves cognitive function in APP/PS1 mice.

**Figure 5.**
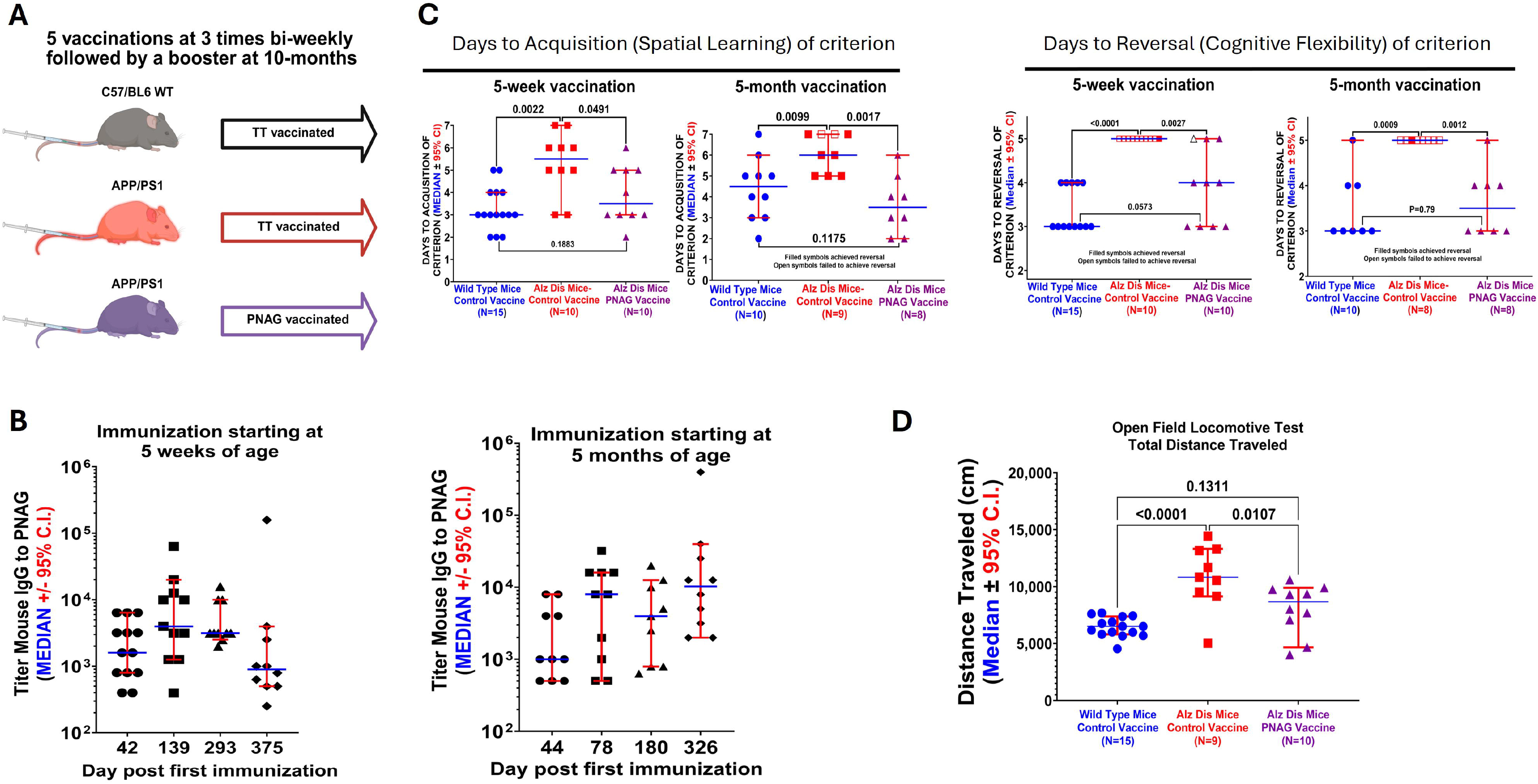
Immune targeting of persistent microbial material preserves cognitive performance in APP/PS1 mice. **(A)** Experimental design for vaccination of wild-type (WT) and APP/PS1 mice with tetanus toxoid (TT) or PNAG–TT beginning at 5 weeks or 5 months of age, followed by booster immunization at 10 months. **(B)** Serum anti-PNAG IgG titers following vaccination. Data are median ± 95% CI. **(C)** Water T-maze performance during acquisition (spatial learning) and reversal (cognitive flexibility). **(D)** Total distance traveled during open-field testing. Data are median ± 95% CI. Statistics: Acquisition Kruskal-Wallis, P = 0.0044 (5-week) and P = 0.0046 (5-month); d*_Cohen_* (APP/PS1-control vs. PNAG vaccine) = 1.261 and 1.549, respectively. Reversal Kruskal-Wallis, P < 0.0001 (5-week) and P = 0.0011 (5-month); d = 2.287 and 2.31, respectively. Open field one-way ANOVA, P < 0.0001; d = 2.244. All large effect sizes. Pairwise comparisons by two-stage linear step-up procedure of Benjamini, Krieger, and Yekutieli (acquisition, reversal) or Tukey’s multiple-comparisons test (open field).

### Immune targeting of persistent microbial material attenuates neuroinflammatory pathways associated with AD pathology

To determine whether the behavioral benefits of PNAG vaccination were accompanied by reduced neuroinflammation, brain tissues from APP/PS1 mice were examined following behavioral testing (*26*). PNAG vaccination markedly attenuated astrocyte and microglial activation compared with TT-vaccinated APP/PS1 controls, with significantly reduced GFAP and IBA1 immunoreactivity (P < 0.0001; Fig. 6A–C) approaching levels observed in non-transgenic mice. Multiplex analysis of whole-brain homogenates detected no significant differences in soluble cytokine concentrations at the experimental endpoint (fig. S4), consistent with the possibility that localized or transient inflammatory responses were not captured by whole-brain measurements. To investigate the underlying mechanism, human THP-1 monocytes expressing GFP-tagged apoptosis-associated speck-like protein containing a CARD (ASC) were exposed to PNAG. PNAG induced robust ASC speck formation, demonstrating inflammasome assembly (Fig. 6D), together with increased secretion of bioactive IL-1β (P < 0.0001; Fig. 6E). Broader cytokine profiling demonstrated increased production of VEGF-α, TNF-α, MCP-1, MCP-2, IL-1β, I-309, GRO-α, and GRO, whereas angiogenin was reduced and IL-10 increased (Fig. 6F). Pharmacologic inhibition of TLR2 significantly reduced IL-1β production, while the pan-caspase inhibitor z-VAD-fmk abolished cytokine release (Fig. 6G), demonstrating that PNAG-induced inflammasome activation requires both TLR2 signaling and caspase activation. Together, these findings demonstrate that immune targeting of persistent microbial material attenuates chronic glial activation in vivo and identify TLR2-dependent inflammasome activation as a mechanism by which persistent microbial material sustains innate immune signaling.

**Figure 6.**
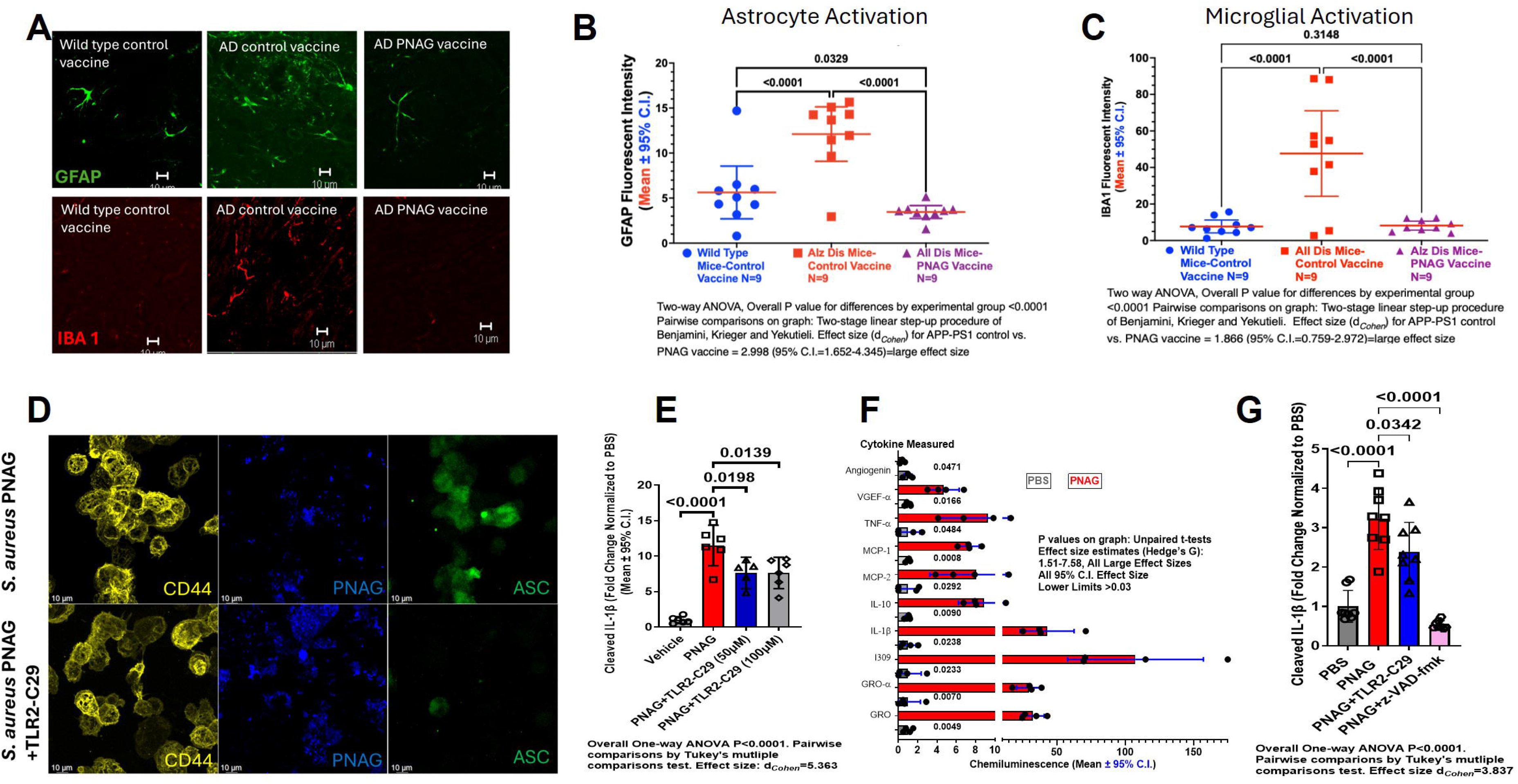
Immune targeting of persistent microbial material attenuates neuroinflammatory responses. **(A)** Representative confocal images of hippocampal sections from wild-type and APP/PS1 mice vaccinated with tetanus toxoid (TT) or PNAG vaccine and stained for GFAP (green) and IBA1 (red). Scale bars, 10 μm. **(B,C)** Quantification of GFAP and IBA1 immunoreactivity. Data are median ± 95% CI. **(D)** Representative confocal images of THP-1 ASC reporter cells exposed to PNAG showing ASC speck formation. Cells were stained for CD44 (yellow), PNAG (blue), and ASC (green). Scale bars, 10 μm. **(E)** Bioactive IL-1β secretion following PNAG exposure. **(F)** Cytokine profiling of THP-1 cells following PNAG exposure. **(G)** Effects of TLR2 inhibition and pan-caspase inhibition on PNAG-induced IL-1β production. Data are mean ± 95% CI.

### Immune targeting of persistent microbial material remodels AD-associated amyloid pathology in APP/PS1 mice

To determine whether immune targeting of persistent microbial material modified AD neuropathology, brain tissue from APP/PS1 mice vaccinated beginning at either 5 weeks or 5 months of age were examined following behavioral testing. PNAG vaccination markedly reduced both plaque-associated PNAG immunoreactivity and amyloid pathology throughout the cortex and hippocampus (Fig. 7A). Compared with TT-vaccinated APP/PS1 mice, vaccinated animals exhibited fewer, smaller, and less compact Aβ plaques together with markedly reduced PNAG–Aβ colocalization. Quantitative analysis confirmed significant reductions in total Aβ immunoreactivity in both the early- and delayed-vaccination cohorts (P < 0.0001 and P = 0.0004, respectively; Fig. 7B). Vaccination also altered amyloid processing. PNAG-vaccinated mice uniquely exhibited detectable soluble Aβ1-38 (P < 0.0001; Fig. 7C), whereas this species was largely absent from TT-vaccinated controls. Insoluble Aβ1-42 was reduced following vaccination, while modest increases in soluble Aβ1-42 were observed in the early-vaccination cohort (fig. S5), resulting in a significantly reduced Aβ1-42:Aβ1-38 ratio (P = 0.0165; Fig. 7C)(*27,28*). To determine whether persistent microbial material similarly influenced amyloid processing in human neurons, Aβ40 and Aβ42 were quantified in conditioned media from PNAG-treated neuronal cultures. PNAG exposure significantly reduced Aβ40 secretion, resulting in an increased Aβ42:Aβ40 ratio relative to vehicle-treated cultures (Fig. 7D). Together, these findings demonstrate that immune targeting of persistent microbial material reduces plaque-associated PNAG, attenuates amyloid deposition, and remodels amyloid processing in APP/PS1 mice. The complementary human neuronal studies further indicate that persistent microbial material directly influences Aβ isoform processing across experimental systems.

**Figure 7.**
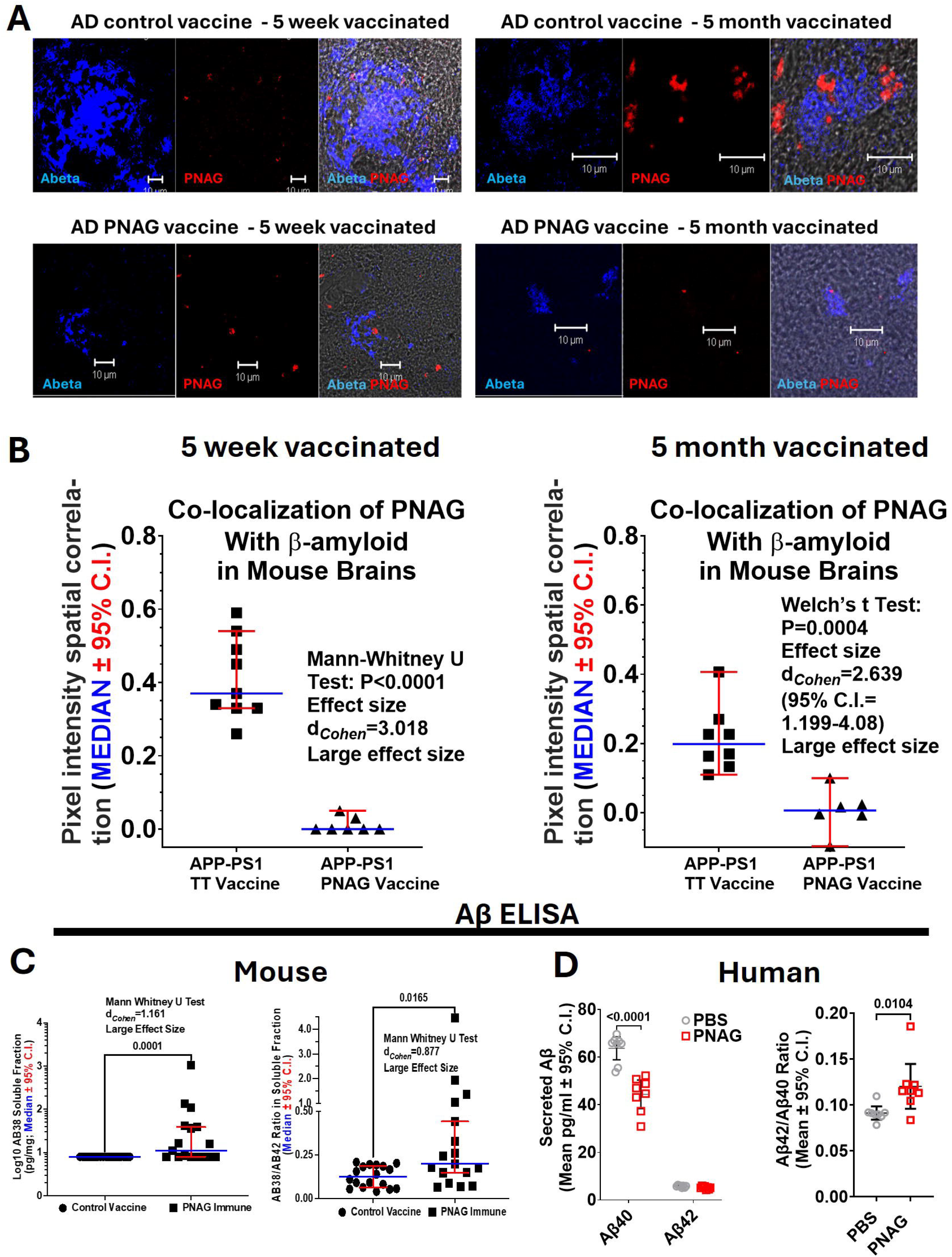
Immune targeting of persistent microbial material remodels amyloid pathology and Aβ processing in APP/PS1 mice. **(A)** Representative confocal images of brain sections from APP/PS1 mice vaccinated with tetanus toxoid (TT) or PNAG vaccine and stained for amyloid-β (Aβ, blue) and PNAG-containing microbial material (red). Scale bars, 10 μm. **(B)** Quantification of spatial colocalization between PNAG-containing microbial material and Aβ in APP/PS1 mice. Data are median ± 95% CI. **(C)** Quantification of soluble Aβ1-38 and the Aβ1-42:Aβ1-38 ratio in brain homogenates. Data are median ± 95% CI. Mann–Whitney *U* test. **(D)** Quantification of Aβ40 and Aβ42 secretion by human neuronal cultures following PNAG exposure. Data are mean ± 95% CI. Statistics: Ordinary two-way ANOVA, P < 0.0001; two-stage linear step-up procedure of Benjamini, Krieger, and Yekutieli multiple-comparisons test (Aβ40: PBS vs. PNAG, P < 0.0001). d*_Cohen_* = 1.548 (95% CI, 0.4389–2.658), large effect size. Aβ42:Aβ40 ratio: Mann–Whitney U test, P = 0.0104; d*_Cohen_* = 1.623, large effect size.

### PNAG vaccination preserves the gut microbial community

One potential explanation for the protective effects of PNAG vaccination is that immunization indirectly alters the intestinal microbiota. To test this possibility, we performed longitudinal 16S rRNA sequencing of fecal samples collected before vaccination and at multiple time points throughout the study (*29*).

Family-level taxonomic profiles remained highly similar between vaccinated and control animals throughout the study (fig. S6A). Measures of alpha and beta diversity demonstrated no significant effects of vaccination on overall microbial community structure (fig. S6B–E). Instead, the modest differences observed reflected genotype, study duration, and cage effects rather than vaccination status. Likewise, differential abundance analysis identified bacterial taxa that differed between APP/PS1 and wild-type mice but were unaffected by PNAG vaccination in APP/PS1 mice (fig. S6F,G). These findings indicate that the therapeutic effects of PNAG vaccination occur without detectable restructuring of the gut microbial community, supporting immune targeting of persistent microbial material rather than alteration of the intestinal microbiota as the mechanism underlying disease modification.

### Implications for Alzheimer disease pathogenesis and therapy

Microbial contributions to AD have attracted increasing attention, but the identity of conserved microbial components capable of sustaining chronic neuroinflammation has remained unclear (*4–9*). Here we identify PNAG-containing microbial material as a conserved component of persistent microbial material associated with human AD pathology and demonstrate that immune targeting of this material modifies disease-relevant outcomes across human tissues, fully human experimental systems, and a murine model of AD. PNAG-containing microbial material colocalized with amyloid plaques in human AD brain, activated TLR2-dependent innate immune signaling, promoted Aβ accumulation and tau phosphorylation in human neuronal systems, and its immune targeting reduced glial activation, amyloid pathology, and cognitive decline in APP/PS1 mice without altering gut microbial community structure.

These findings extend the antimicrobial protection hypothesis (*7,8*), which proposes that Aβ functions as an innate immune effector produced in response to microbial exposure. Within this framework, Aβ deposition may initially limit dissemination of microbial structures. Our findings suggest that persistent microbial material may provide a sustained inflammatory stimulus that converts an initially protective response into chronic neuroinflammation. The accumulation of PNAG-containing microbial material within plaque-associated regions of human AD brain, together with the ability of purified PNAG and PNAG-containing microbial vesicles to induce AD-associated pathology in fully human neuronal systems, identifies persistent microbial material as a biologically active component of this process and establishes a tractable experimental target for therapeutic intervention.

The vaccination studies further demonstrate that immune targeting of naturally encountered microbial material is sufficient to modify disease progression in the absence of experimental infection or exogenous PNAG challenge. Animals were exposed only to their endogenous microbiota and routine environmental microbial exposures, yet vaccination improved cognition, reduced chronic glial activation, attenuated amyloid pathology, and shifted amyloid processing toward increased soluble Aβ1-38 and a reduced Aβ1-42:Aβ1-38 ratio. Because shorter Aβ isoforms exhibit reduced aggregation propensity (*26–28*), these findings suggest that reducing the burden of persistent microbial material remodels amyloid processing indirectly through upstream inflammatory mechanisms rather than by directly targeting Aβ itself.

The fully human mechanistic studies provide a potential explanation for these observations. Across neuronal monocultures, three-dimensional neuron–astrocyte–microglia models, and human THP-1 monocytes, persistent microbial material consistently activated TLR2-dependent inflammasome signaling, increased bioactive IL-1β production, and promoted accumulation of AD-associated proteins. These responses occurred in the absence of viable microorganisms, demonstrating that microbial material alone is sufficient to sustain innate immune activation. Although additional pattern-recognition receptors and downstream signaling pathways likely contribute to AD pathology, these findings identify TLR2-dependent recognition as one experimentally defined mechanism linking persistent microbial material to chronic neuroinflammation.

An important implication of this work is that immune targeting of persistent microbial material differs fundamentally from approaches aimed at eliminating microorganisms or broadly restructuring the intestinal microbiota. PNAG vaccination produced no detectable changes in gut microbial community composition, indicating that therapeutic benefit was not mediated through microbiome disruption but rather through selective targeting of persistent microbial material while preserving the host microbial ecosystem. This strategy may therefore be applicable to chronic inflammatory diseases in which persistent microbial material continues to stimulate innate immunity despite the absence of ongoing infection.

This study has several limitations. We did not establish the anatomical origin or trafficking pathways of persistent microbial material within the human brain, nor do we propose that PNAG is the only microbial component contributing to chronic neuroinflammation. Rather, PNAG serves as an experimentally tractable representative of a broader class of conserved microbial material capable of sustaining innate immune activation. Future studies should define the sources, trafficking mechanisms, and clearance pathways of persistent microbial material and determine whether therapeutic targeting of this process translates to human disease.

Together, these findings identify persistent microbial material as a therapeutically targetable upstream driver of chronic neuroinflammation that contributes to AD pathology. More broadly, they establish a framework in which conserved microbial material functions not simply as a marker of prior microbial exposure but as a biologically active and therapeutically modifiable contributor to chronic inflammatory disease.

## Acknowledgements

**General:** We thank our technical support teams and the Metabolism and Mitochondrial Research Core (Beth Isreal Deaconess Medical Center, Boston, MA) Laboratory Research – Center for Resuscitation Science for their contributions. We thank Dr. Kimberly Jefferson of Virginia Commonwealth University School of Medicine for provision of the E. coli strains.

## Funding

Infectious Diseases Society of America (IDSA)(C.C.B)

Multidisciplinary University Research Initiatives (MURI) W911NF-23-1-0276 (C.C.B.,D.L.K)

## Author Contributions

Conceptualization: CCB, GBP, ECM

Methodology: CCB, GBP, ECM, BC, CAL

Investigation CCB, GBP, ECM, BC, MV, TZ, NN, AA, GS, DL, EL, AA, LHP, CR, EC, CW

DC, MFK, LMC, CAL

Visualization: CCB, ECM, GBP

Funding acquisition: CCB, DLK

Project administration: CCB, ECM,GBP, DLK

Supervision: CCB, ECM, GBP, DLK

Writing - original draft: CCB, ECM, GBP

Writing - review & editing: CCB, ECM, GBP, DC, DLK, CAL

## Data and materials availability

All data supporting the findings of this study are provided in the main text or the Supplementary Materials. Raw datasets and analysis code are available from the corresponding author upon reasonable request.

## Supplementary Materials

## Materials and Methods

### Study Design

This study was designed to evaluate the contribution of PNAG-containing microbial material to AD pathology and to assess the therapeutic potential of PNAG-targeted vaccination. The experimental design integrated human observational studies, fully human mechanistic models, and in vivo preclinical testing in APP/PS1 mice to align mechanistic, pathological, biomarker, and functional endpoints across complementary experimental systems. Postmortem brain tissue from patients with AD and age- and sex-matched controls was analyzed to determine the spatial relationship between PNAG-containing microbial material and Aβ pathology. Mechanistic studies were performed using two-dimensional and three-dimensional human induced neural stem cell (hiNSC)-derived neuronal cultures, neuron–astrocyte–microglia tri-cultures (*21,22,24*), and human THP-1 monocytes to determine whether PNAG-containing microbial material activates inflammasome signaling and promotes accumulation of AD-associated proteins. Pharmacologic inhibition of Toll-like receptor 2 (TLR2) together with antibody-mediated neutralization of PNAG were used to investigate underlying mechanisms. Preclinical studies evaluated whether vaccination against PNAG modifies cognitive, pathological, and biomarker outcomes in APP/PS1 mice, including behavioral performance, glial activation (IBA1 and GFAP), and Aβ isoform profiles. Animals were randomized to receive either a PNAG–tetanus toxoid (PNAG-TT) conjugate vaccine or tetanus toxoid (TT) alone (*23*), with investigators blinded during behavioral testing, tissue analysis, and data interpretation. TT alone was selected as the control because it constitutes the carrier protein for the conjugate vaccine, constitutes 85% of the vaccine by weight, is itself a widely used vaccine in humans and therefore effects from high level immunity induced by the TT component due to multiple immunizations in mice would control for any carrier effects mimicking the human response to this antigen. Overall, this strategy controls for adjuvant, carrier protein, immunization schedule, the majority of vaccine mass and a mimic of human immune responses to the current PNAG vaccine construct, while isolating immune responses directed against the PNAG glycan. Sample sizes were based on preliminary studies to provide adequate power for molecular and behavioral endpoints. Longitudinal 16S rRNA sequencing of fecal samples was performed to determine whether vaccination altered gut microbial community composition.

### Human postmortem brain tissue

Postmortem brain tissue from patients with AD and age- and sex-matched control individuals without a diagnosis of neuroinflammatory disease at death was obtained from the Massachusetts General Hospital AD Research Center (MGH ADRC) following institutional ethical approval. Sections were stained with a fully human IgG1 monoclonal antibody specific for poly-*N*-acetylglucosamine (PNAG; MAb F598)(*23,16*) together with an anti-Aβ antibody (6E10; BioLegend, SIG-39320). Isotype control staining was performed using MAb F429 (*16*). To assess glial activation, sections were stained with antibodies against ionized calcium-binding adaptor molecule 1 (IBA1; Wako, 019-19741) and glial fibrillary acidic protein (GFAP; Sigma, G3893). Images were acquired using a Zeiss LSM confocal microscope (Carl Zeiss, Oberkochen, Germany) and analyzed for spatial colocalization and mean fluorescence intensity (MFI) using Axiovert software.

### Experimental mice

Male APP/PS1 transgenic mice (B6C3-Tg(APPswe,PSEN1dE9)85Dbo/Mmjax) and non-transgenic male littermates were maintained under specific pathogen-free conditions. Mice were immunized subcutaneously with 30 μg of PNAG oligosaccharide (5GlcNH₂) conjugated to tetanus toxoid (TT; Alopexx, Inc., Cambridge, MA) formulated in 0.2% Alhydrogel (InvivoGen, vac-alu-50) in a total volume of 200 μl (*23*). Control animals received an identical formulation containing tetanus toxoid and adjuvant but lacking the PNAG oligosaccharide. Immunizations were administered at 5, 8, 11, and 14 weeks of age and every three months thereafter until euthanasia. Blood was obtained periodically over this time period to assess immune responses to native PNAG antigen by ELISA. Each experimental cohort initially consisted of 15 male mice but over the course of the experimental period some were lost primarily due to injuries from fighting with cage mates so final group sizes were 8-11 mice.

### PNAG-specific antibody titers

Serum IgG titers to PNAG were quantified by indirect enzyme-linked immunosorbent assay (ELISA). Microtiter plates were coated with purified PNAG isolated from *Acinetobacter baumannii* (*16,18*). Bound antibodies were detected using alkaline phosphatase-conjugated anti-mouse IgG (Jackson ImmunoResearch, 115-055-174) and 4-nitrophenyl phosphate disodium salt hexahydrate substrate (Sigma, N4645).

### Murine behavioral assessment

Behavioral testing consisted of a 30-minute open-field locomotor assay (OFL; automated tracking using Noldus EthoVision) and the water T-maze (WTM) to assess spatial acquisition and reversal learning (10 trials per day). Behavioral testing was performed between 12 and 15 months of age.

### Mouse brain Aβ isoform and cytokine analysis

Brain tissues collected following completion of behavioral testing were homogenized in RIPA buffer containing protease and phosphatase inhibitors (Roche, 04693116001 and C764L25). Soluble and insoluble fractions were separated by centrifugation. Aβ1-38, Aβ1-40, Aβ1-42, and cytokine concentrations were quantified using Meso Scale Discovery (MSD) multiplex assays (V-PLEX Panel 1, 4G8: mouse and human Aβ; Mouse Proinflammatory Panel 1) according to the manufacturer’s instructions. Total protein concentrations were determined using a bicinchoninic acid (BCA) assay (Pierce, 23225).

### HEK-Blue reporter assay for bioactive IL-1β

HEK-Blue IL-1β reporter cells (InvivoGen, hkb-il1bv2) were maintained according to the manufacturer’s instructions in high-glucose DMEM (Gibco, 11965092) supplemented with 10% heat-inactivated fetal bovine serum (Gibco, A5256801) and a 1% antibiotic-antimycotic reagent (Gibco, 15240062). Conditioned media collected from experimental cultures were transferred to HEK-Blue IL-1β reporter cells and incubated overnight. In this reporter system, biologically active cleaved IL-1β stimulates the stably expressed IL-1 receptor, inducing secretion of embryonic alkaline phosphatase (SEAP), thereby providing a functional readout of bioactive IL-1β. SEAP activity was quantified using QUANTI-Blue reagent (InvivoGen, rep-qbs), and absorbance was measured at 630 nm according to the manufacturer’s instructions.

### Cell culture

Human induced neural stem cells (hiNSCs) were generated by direct lineage reprogramming of neonatal foreskin fibroblasts as previously described (*21,22*). hiNSCs were maintained in KnockOut DMEM supplemented with KnockOut Serum Replacement XenoFree Medium, GlutaMAX, recombinant human basic fibroblast growth factor (bFGF), β-mercaptoethanol, and antibiotic-antimycotic (all Gibco) under standard culture conditions. Human microglia clone 3 cells (HMC3; ATCC, CRL-3304) were maintained in RPMI 1640 supplemented with 10% heat-inactivated fetal bovine serum (HI-FBS) and 1% antibiotic-antimycotic. Primary human astrocytes (ScienCell, 1800) were cultured in complete astrocyte medium (ScienCell, 1801) according to the manufacturer’s recommendations. THP-1 human monocytes were maintained in RPMI 1640 supplemented with 10% HI-FBS and 1% antibiotic-antimycotic and differentiated using phorbol 12-myristate 13-acetate (PMA) before experimentation. Human NGN2-induced pluripotent stem cells (iPSCs) were generated using a PiggyBac transposon system and differentiated into neurons as previously described (*24*). Mature neurons were maintained in BrainPhys medium supplemented with GlutaMAX, B-27, and antibiotic-antimycotic prior to experimental treatment.

### Purification of PNAG for cellular studies

PNAG was purified from *Acinetobacter baumannii* strain S1 as previously described (*13–19*) or from the PNAG-overproducing *Staphylococcus aureus* strain MN8M. To purify PNAG from *S. aureus*, cultures of strain MN8M were grown for 36 h in RPMI medium, and cell-free supernatants were subjected to ethanol precipitation. The resulting insoluble material was sequentially treated with DNase I, RNase A, and proteinase K before recovery by centrifugation of the PNAG-containing fraction, acid solubilization, dialysis first against pH 2.8 0.1M glycine buffer then against water, and lyophilization. Purity was confirmed by 1H nuclear magnetic resonance (NMR) spectroscopy and exceeded 99% (*19*). Purified PNAG was dissolved in 5 N HCl, immediately neutralized with an equal volume of 5 M NaOH, and diluted in sterile PBS or 0.04 M phosphate buffer (pH 7.4) before experimental use.

### Isolation of microbial membrane vesicles

*Escherichia coli* outer membrane vesicles (OMVs) were isolated from overnight cultures of strain UTI-J-derived mutants D*csrA* (PNAG producer) or D*nhrA* (PNAG deficient)(*25*). Cell-free culture supernatants were collected by centrifugation, filtered (0.2 μm), and purified using the ExoBacteria OMV Isolation Kit (System Biosciences). *Staphylococcus aureus* bacterial extracellular vesicles (BEVs) were isolated from the PNAG-overproducing strain MN8M or the PNAG-deficient mutant MN8 Δ*ica*. Cell-free culture supernatants were filtered (0.2 μm), concentrated using 100-kDa Centricon® Plus-70 centrifugal filter units (Merck, UFC710008), and ultracentrifuged at 100,000 × *g* for 3 h at 4°C. Vesicle pellets were resuspended in PBS.

Protein concentrations of OMV and BEV preparations were determined using a bicinchoninic acid (BCA) assay before use in subsequent experiments.

### 2D in vitro PNAG treatment

hiNSCs were differentiated for 7 days in BrainPhys medium supplemented with 1% antibiotic-antimycotic (Gibco, 15240062), 1% GlutaMAX (Gibco, 35050061), and 2% B-27 (Gibco, 17504044) following plating onto poly-L-ornithine/laminin-coated culture plates. Differentiated cultures were treated with purified *S. aureus* or *A. baumannii* PNAG (10 μg/mL), *E. coli* OMVs (5 μg/mL), *S. aureus* BEVs (5 μg/mL), or PBS control for 4 or 7 days without media replacement.

THP-1 monocytes were differentiated with phorbol 12-myristate 13-acetate (PMA) before treatment with purified PNAG (10 μg/mL), *E. coli* OMVs (5 μg/mL), *S. aureus* BEVs (5 μg/mL), or PBS for 24–48 h.

### Scaffold fabrication

Silk fibroin (6% w/v) scaffolds were fabricated using established methods as previously described (*21,22,24*). Briefly, porous silk scaffolds were generated by salt leaching, stabilized by β-sheet formation, extensively washed to remove residual salt, and cored into 3-mm diameter × 1-mm height scaffolds. Scaffolds were sterilized by autoclaving and stored at 4°C until use.

### 3D human brain-like cultures

Three-dimensional neuronal monocultures and neuron–astrocyte–microglia tri-cultures were generated using porous silk fibroin scaffolds as previously described (*21,22,24*). Laminin-coated scaffolds were seeded with 1 × 10⁶ human neurons alone or with 1 × 10⁶ neurons, 5 × 10⁵ astrocytes, and 2.5 × 10⁵ microglia before embedding in type I collagen hydrogel. Cultures were maintained for 4 weeks with twice-weekly half-medium changes to permit tissue maturation before experimental treatment. Mature cultures were treated with purified *S. aureus* PNAG (10 μg/mL) or PNAG-positive or PNAG-deficient *E. coli* outer membrane vesicles (OMVs; 5 μg/mL) for 4–10 days before downstream analyses (*25*).

### Immunofluorescence microscopy

Two-dimensional cultures and three-dimensional scaffold cultures were fixed with 4% paraformaldehyde, permeabilized, blocked, and stained using standard immunofluorescence procedures. Primary antibodies were incubated overnight at 4°C, followed by fluorophore-conjugated secondary antibodies. Antibodies used for each experiment are listed in **table S1**.

Confocal imaging was performed using a Leica TCS SP8 microscope (Leica Microsystems, Wetzlar, Germany). Z-stack images were acquired using 25× and 40× objectives with identical acquisition settings within each experiment to permit quantitative comparison of fluorescence intensity.

### Molecular analyses of human cell cultures

Conditioned media from neuronal cultures were collected after 4 days of treatment for quantification of human Aβ40 (Invitrogen, KHB3481) and ultrasensitive Aβ42 (Invitrogen, KHB3544) by enzyme-linked immunosorbent assay (ELISA) according to the manufacturer’s instructions. For analysis of neuronal cell lysates following 7 days of treatment, Aβ42 measurements were normalized to 10 μg total protein per well. Phosphorylated tau (pTau; Ser202/Thr205) was quantified in human induced pluripotent stem cell-derived neuronal lysates normalized to 10 μg total protein using the AT8 ELISA kit (RayBiotech, PEL-Tau-S202). Cytokine secretion from THP-1 cultures was assessed after 48 h of treatment using the RayBiotech C-Series Human Cytokine Array (AAH-CYT-3), and relative cytokine abundance was quantified by chemiluminescence according to the manufacturer’s instructions. Total RNA was isolated using the RNeasy Mini Kit (Qiagen, 74104), reverse transcribed using the Bio-Rad Reverse Transcription Supermix (1708841), and analyzed by quantitative RT-PCR using iTaq Universal SYBR Green Supermix on a CFX96 Real-Time PCR Detection System (Bio-Rad). GAPDH served as the reference gene. Primer sequences are listed in **table S2**.

### Microbiota profiling

Fecal pellets were collected from APP/PS1 mice before vaccination and 44, 78, and 400 days after vaccination. Microbial DNA was extracted using the Qiagen PowerSoil PowerLyzer Kit. The V4 region of the bacterial 16S rRNA gene was amplified using primers 515F/806R and sequenced on an Illumina MiSeq platform. Alpha- and beta-diversity analyses were performed using QIIME2 (*29,30*).

### Study approval

All animal procedures were approved by the Brigham and Women’s Hospital and Harvard Medical Area Institutional Animal Care and Use Committees (IACUCs) and were performed in accordance with institutional guidelines.

### Statistical analysis

Data distributions were assessed for normality, log-normality, or non-parametric behavior before statistical testing. Appropriate two-sided parametric or non-parametric tests were selected for pairwise and multiple-group comparisons. Multi-group comparisons were analyzed using one-way ANOVA followed by appropriate post hoc testing or, where data were non-parametric, Kruskal–Wallis analysis with Dunn’s multiple-comparison test. Each data point represents an independent biological replicate (brain, mouse, culture well, or 3D scaffold), and sample sizes (n) are indicated in the corresponding figure panels. Statistical tests, exact *P* values, effect size estimates, and 95% confidence intervals (where applicable) are reported in the figures.

### Supplementary Text

**Figure S1.**
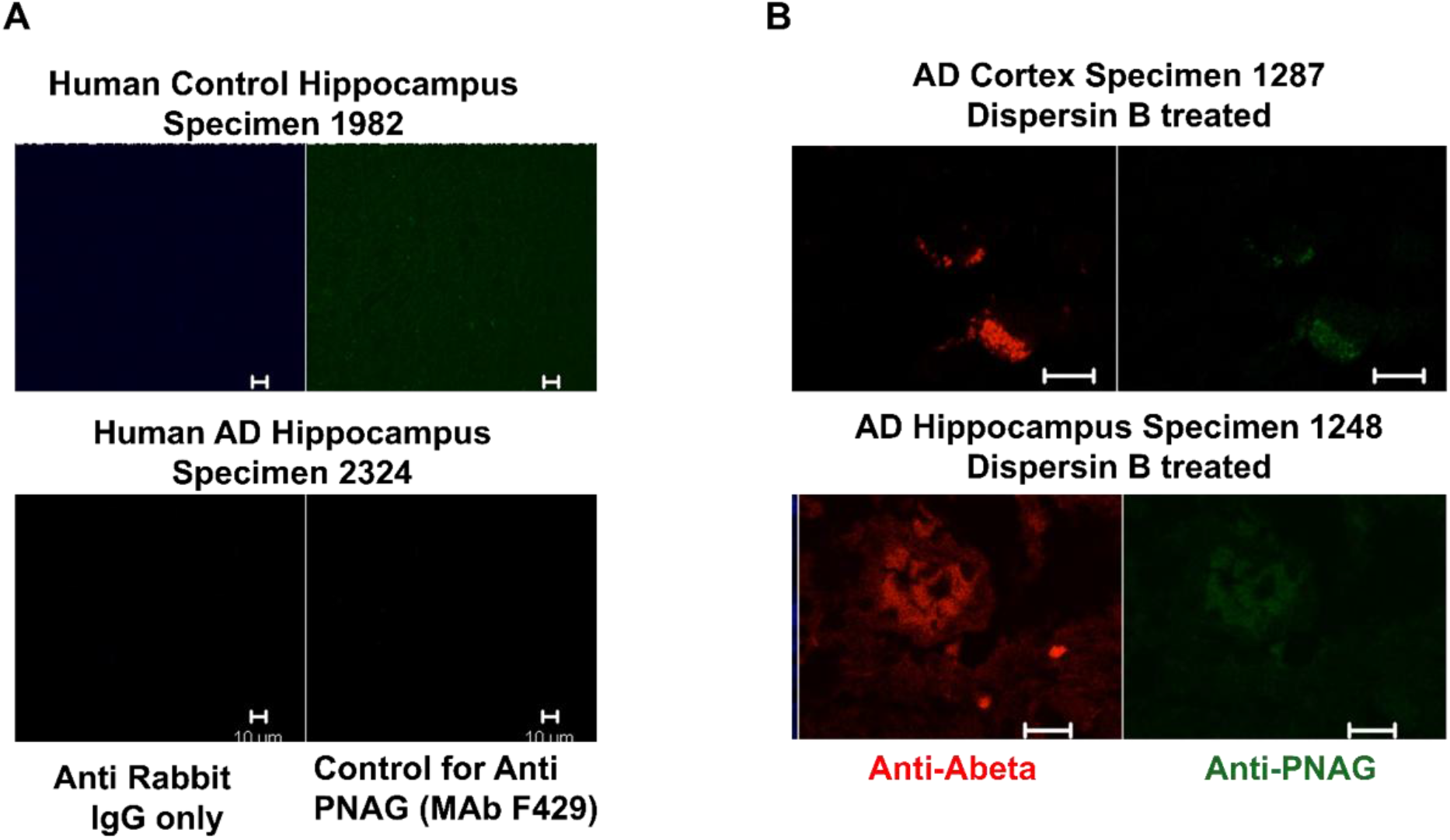
Validation of anti-PNAG immunostaining specificity. **(A)** Negative control staining of postmortem human brain sections using secondary antibody alone or the isotype control monoclonal antibody F429. No specific fluorescence was detected.**(B)** Human Alzheimer disease brain sections before and after treatment with the PNAG-specific glycosidase Dispersin B. Dispersin B abolished anti-PNAG (F598) immunoreactivity while preserving amyloid-β (Aβ) staining. Scale bars, 10 μm.

**Figure S2.**
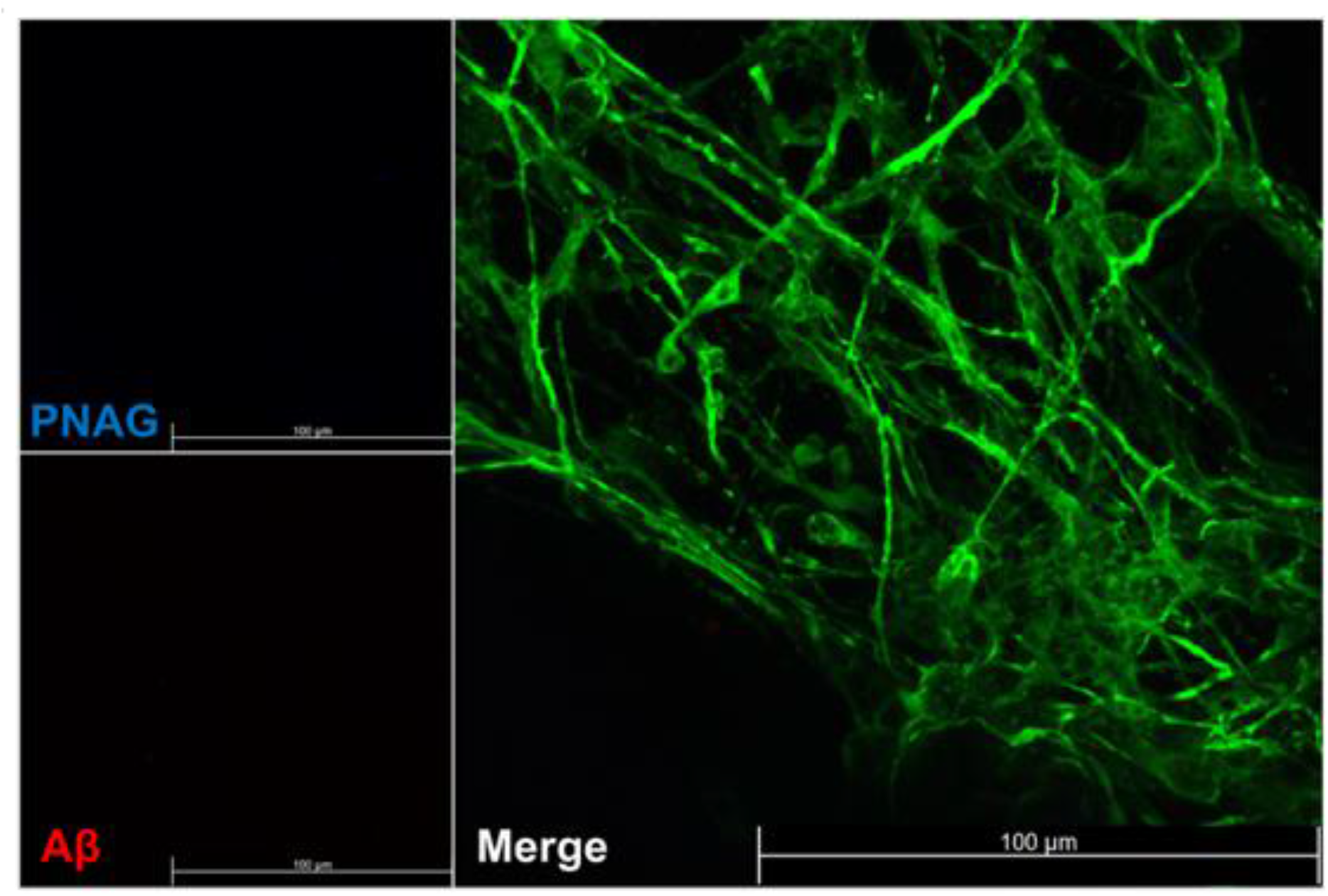
Baseline PNAG and amyloid-β staining in untreated three-dimensional human brain cultures. Representative confocal image of a four-day PBS-treated three-dimensional human neuron astrocyte–microglia (NAM) silk scaffold culture stained for MAP2 (green), PNAG-containing microbial material (blue), and amyloid-β (Aβ, red). Scale bar, 100 μm.

**Figure S3.**
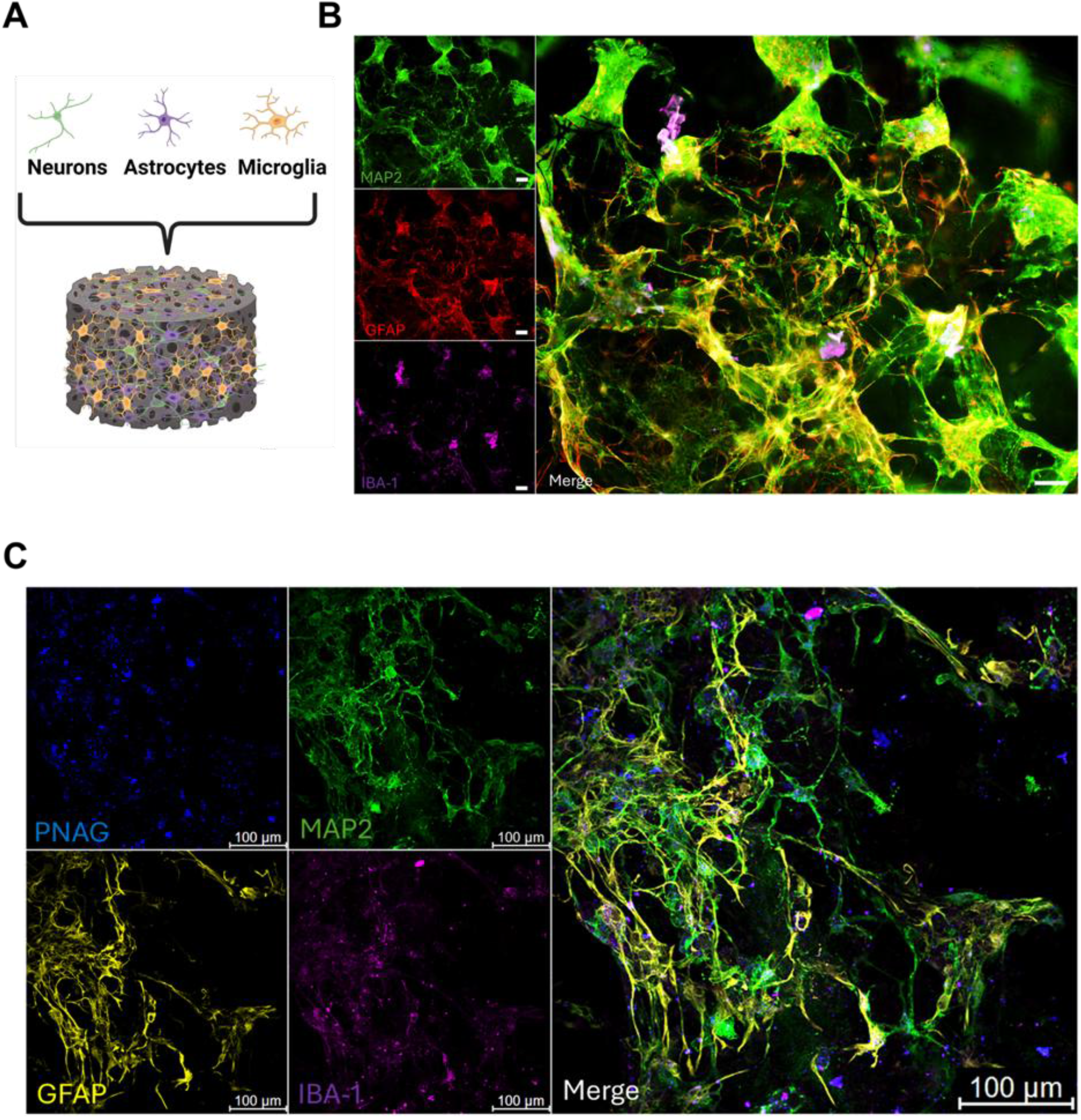
Characterization of the three-dimensional human neuron–astrocyte–microglia (NAM) culture system. **(A)** Schematic of the three-dimensional silk scaffold containing human neurons, astrocytes, and microglia (NAM culture). **(B)** Representative confocal images of mature NAM cultures stained for MAP2 (green), GFAP (red), and IBA1 (magenta). **(C)** Representative confocal images of NAM cultures following 4-day PNAG exposure stained for PNAG-containing microbial material (blue), MAP2 (green), GFAP (yellow), and IBA1 (magenta). Scale bars, 100 μm.

**Figure S4.**
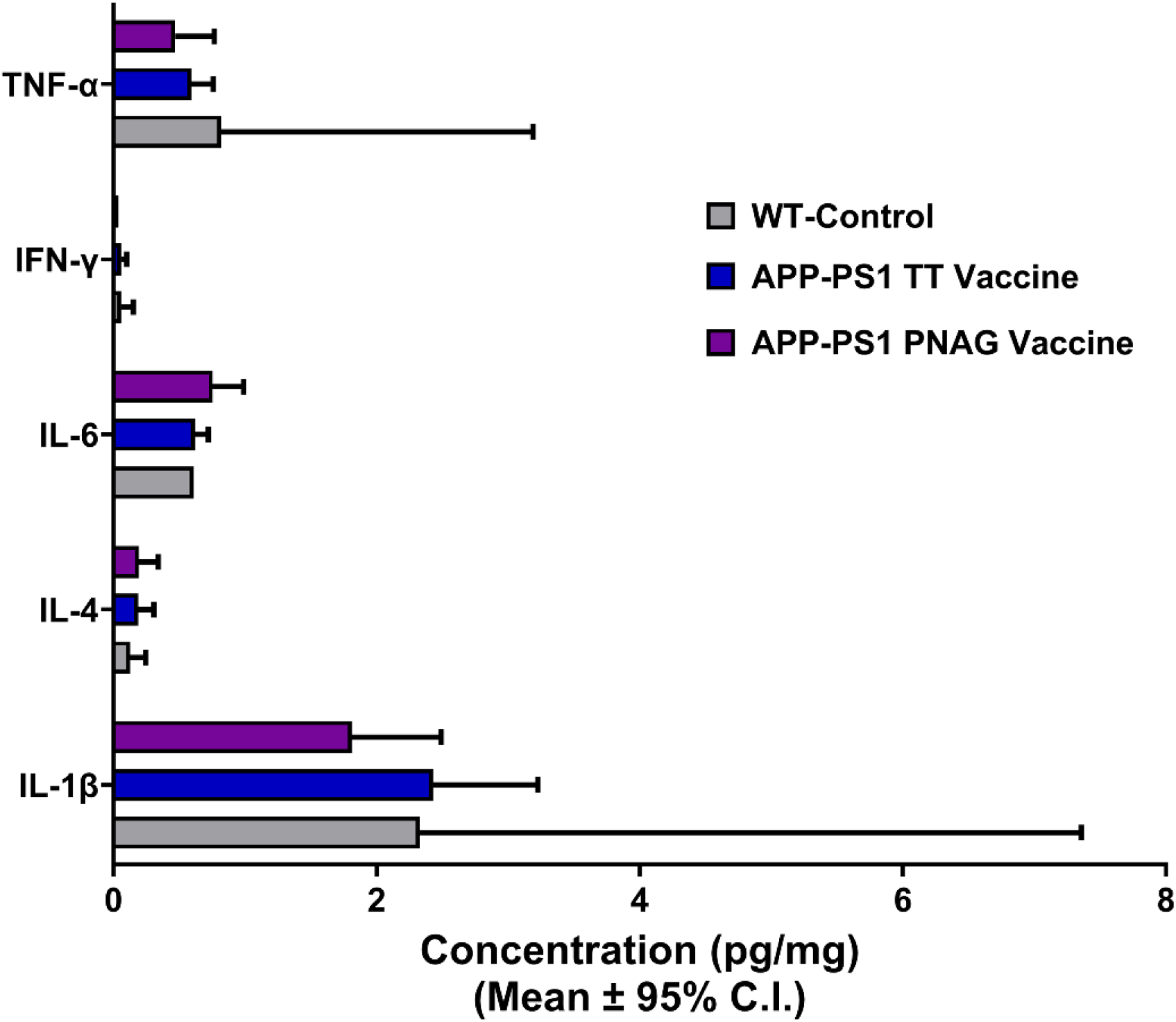
Whole-brain cytokine concentrations following PNAG vaccination. Whole-brain homogenates from 15-month-old APP/PS1 mice vaccinated with tetanus toxoid (TT) or PNAG vaccine were analyzed using a Meso Scale Discovery (MSD) cytokine panel. No significant differences were detected between groups.

**Figure S5.**
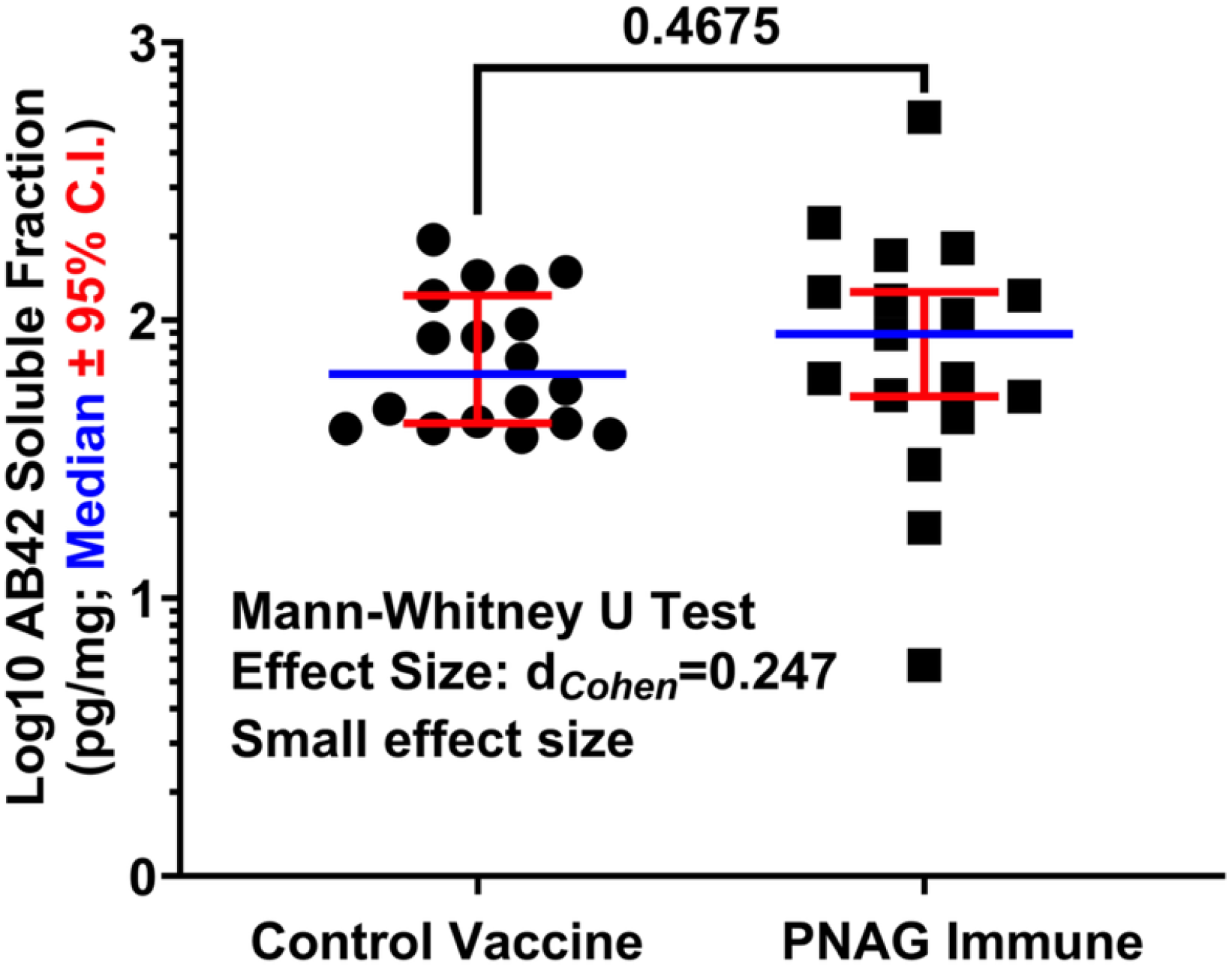
Soluble Aβ42 concentrations following PNAG vaccination. Whole-brain soluble Aβ42 concentrations were quantified in APP/PS1 mice vaccinated with tetanus toxoid (TT) or PNAG vaccine using the Meso Scale Discovery (MSD) V-PLEX assay. Soluble Aβ42 concentrations were modestly increased in PNAG-vaccinated mice. Data are median ± 95% CI.

**Figure S6.**
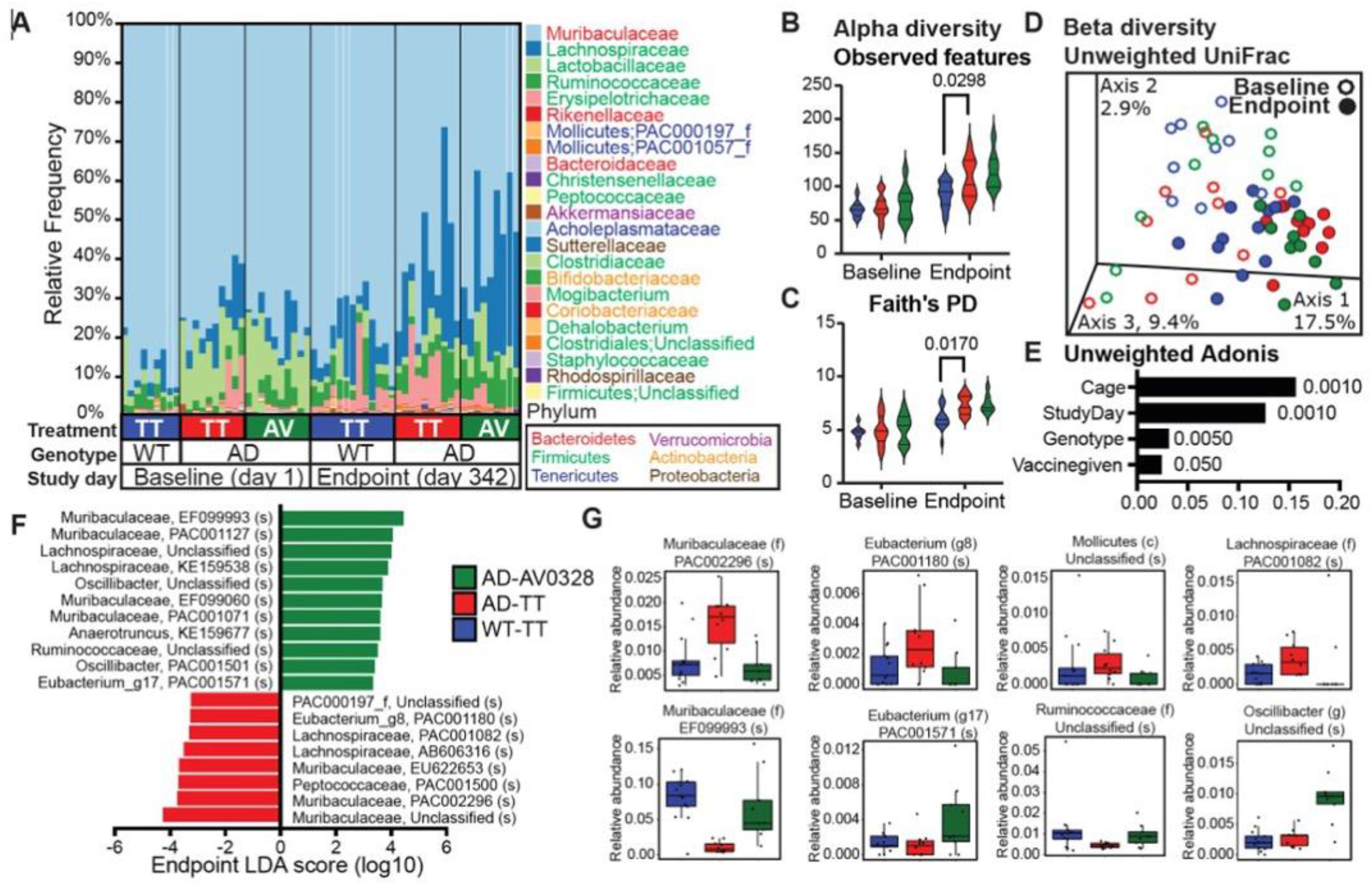
PNAG vaccination does not alter gut microbial community structure. **(A)** Family-level taxonomic composition of fecal microbiota from wild-type (WT) and APP/PS1 (AD) mice vaccinated with tetanus toxoid (TT) or PNAG vaccine at baseline (day 1) and study endpoint (day 342). **(B,C)** Alpha diversity measured by observed features and Faith’s phylogenetic diversity (PD). **(D)** Principal coordinates analysis of unweighted UniFrac distances. **(E)** PERMANOVA (ADONIS) analysis of factors contributing to microbial community variation. **(F,G)** Differential abundance analyses of taxa identified by LEfSe.

**Supplementary Table S1.**
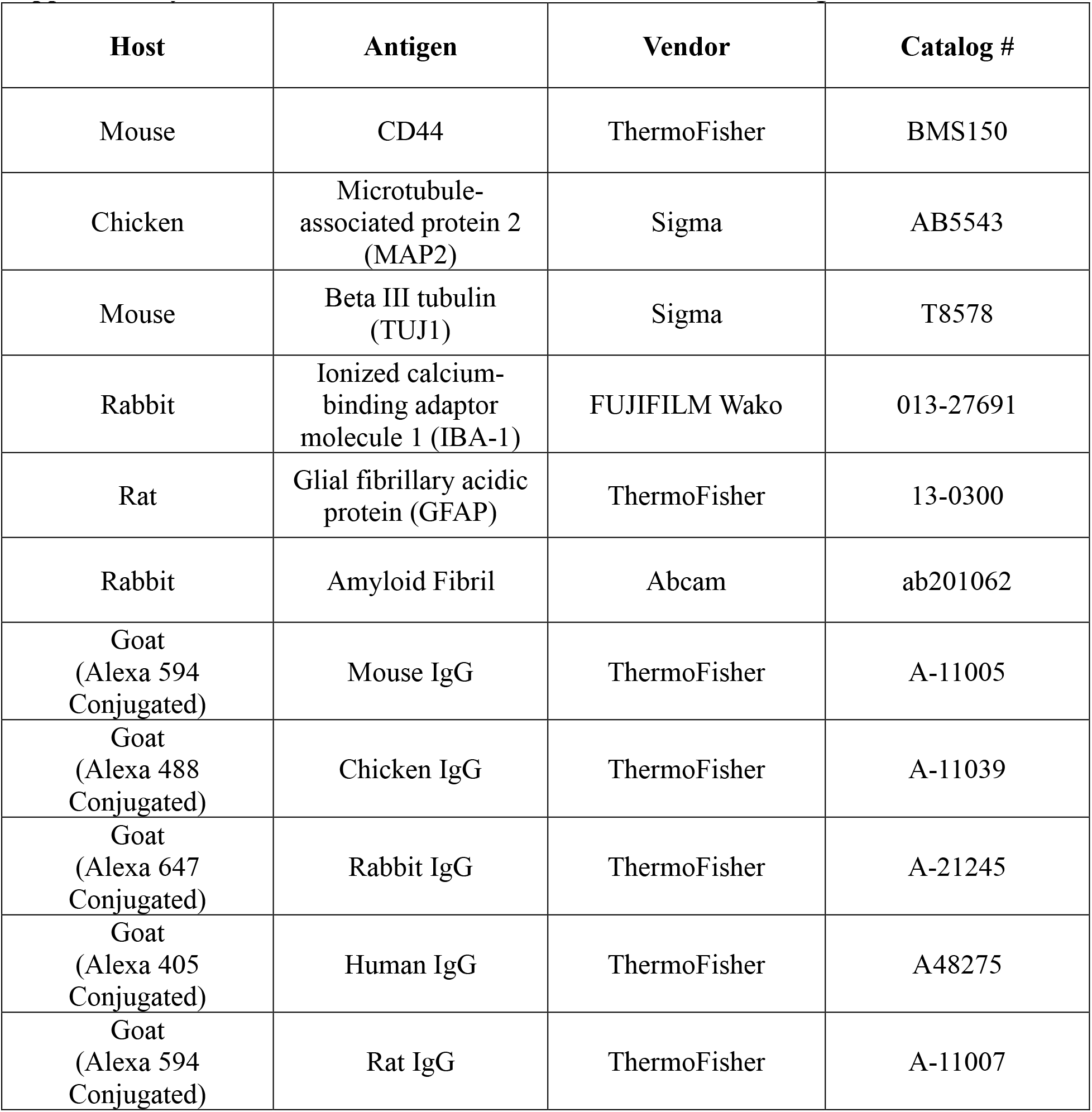
Antibodies used for *in vitro* immunostaining.

**Supplementary Table S2.**
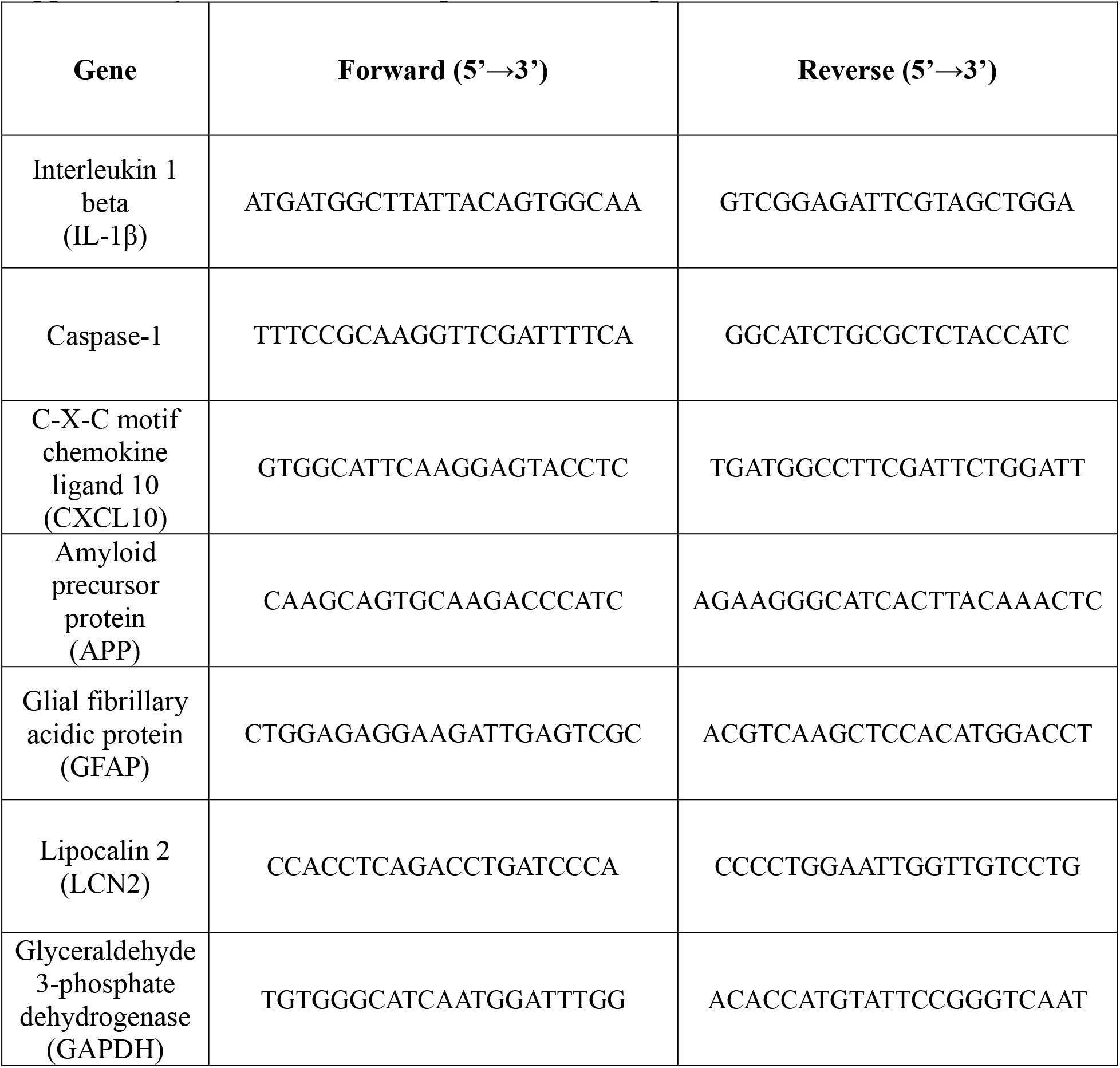
Primer sequences used for qRT-PCR.

