## Supplementary figures and images for "Persistent microbial material contributes to Alzheimer disease and is targetable by vaccination"

### Supplemental Figure S1

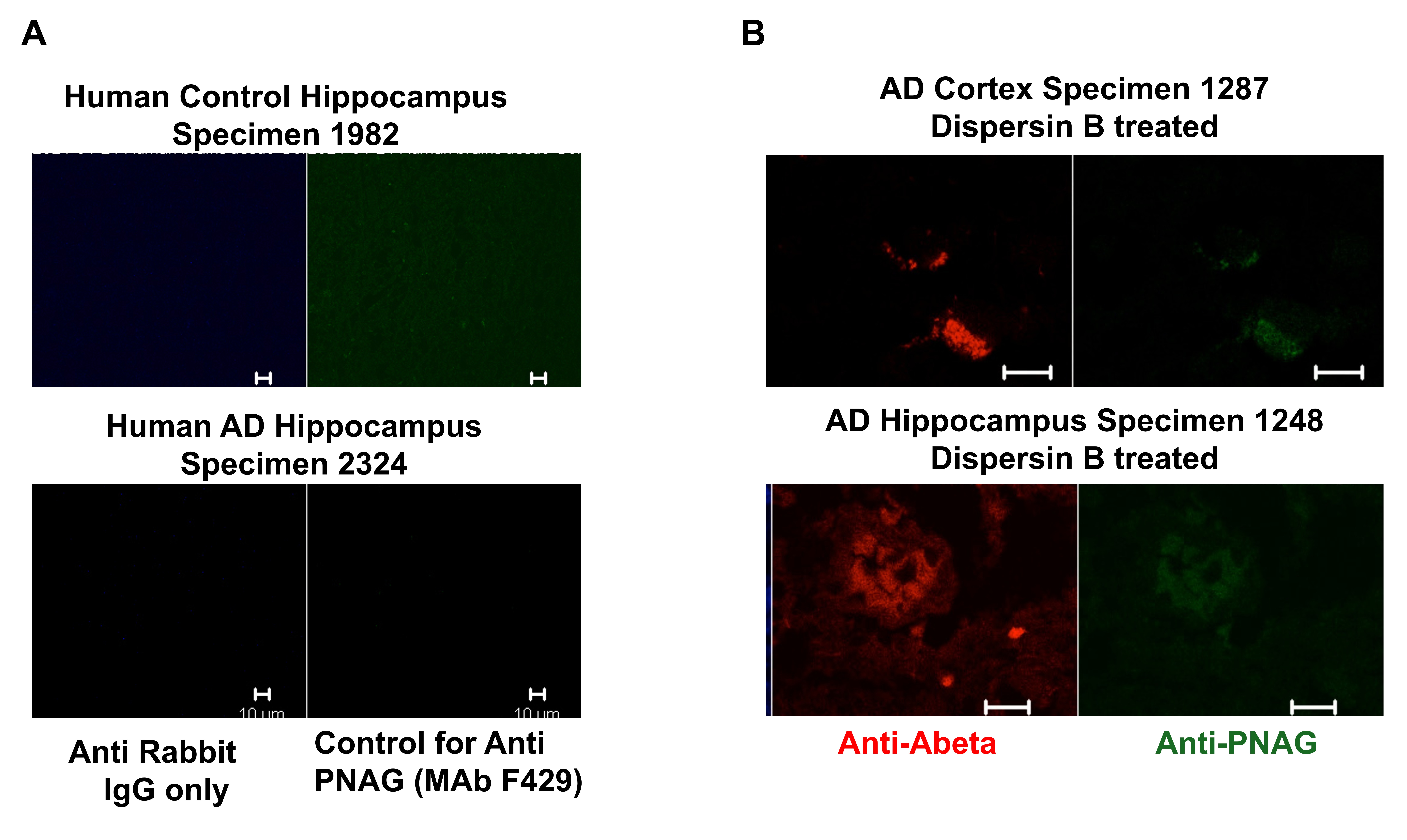

### Supplemental Figure S2

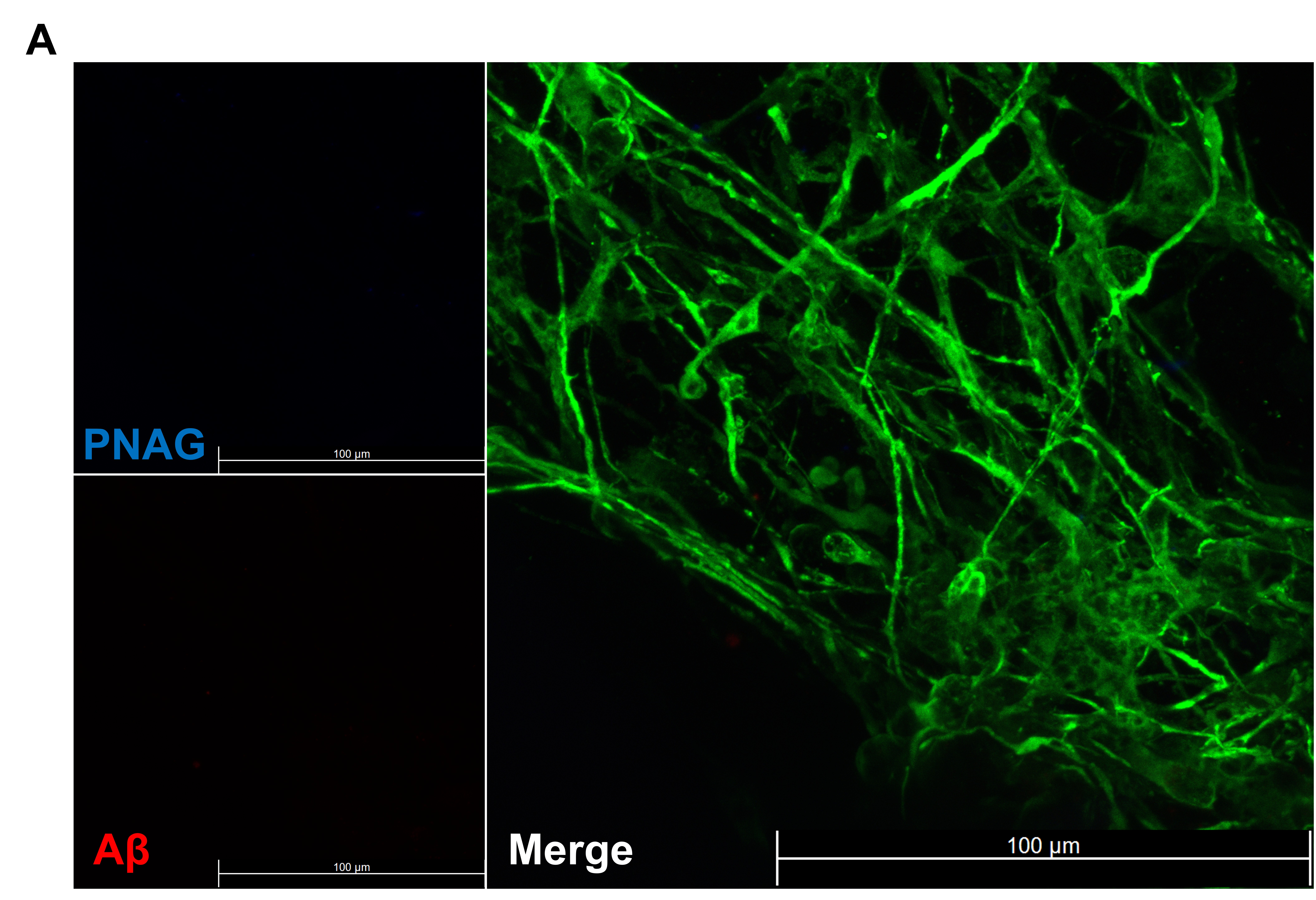

### Supplemental Figure S3

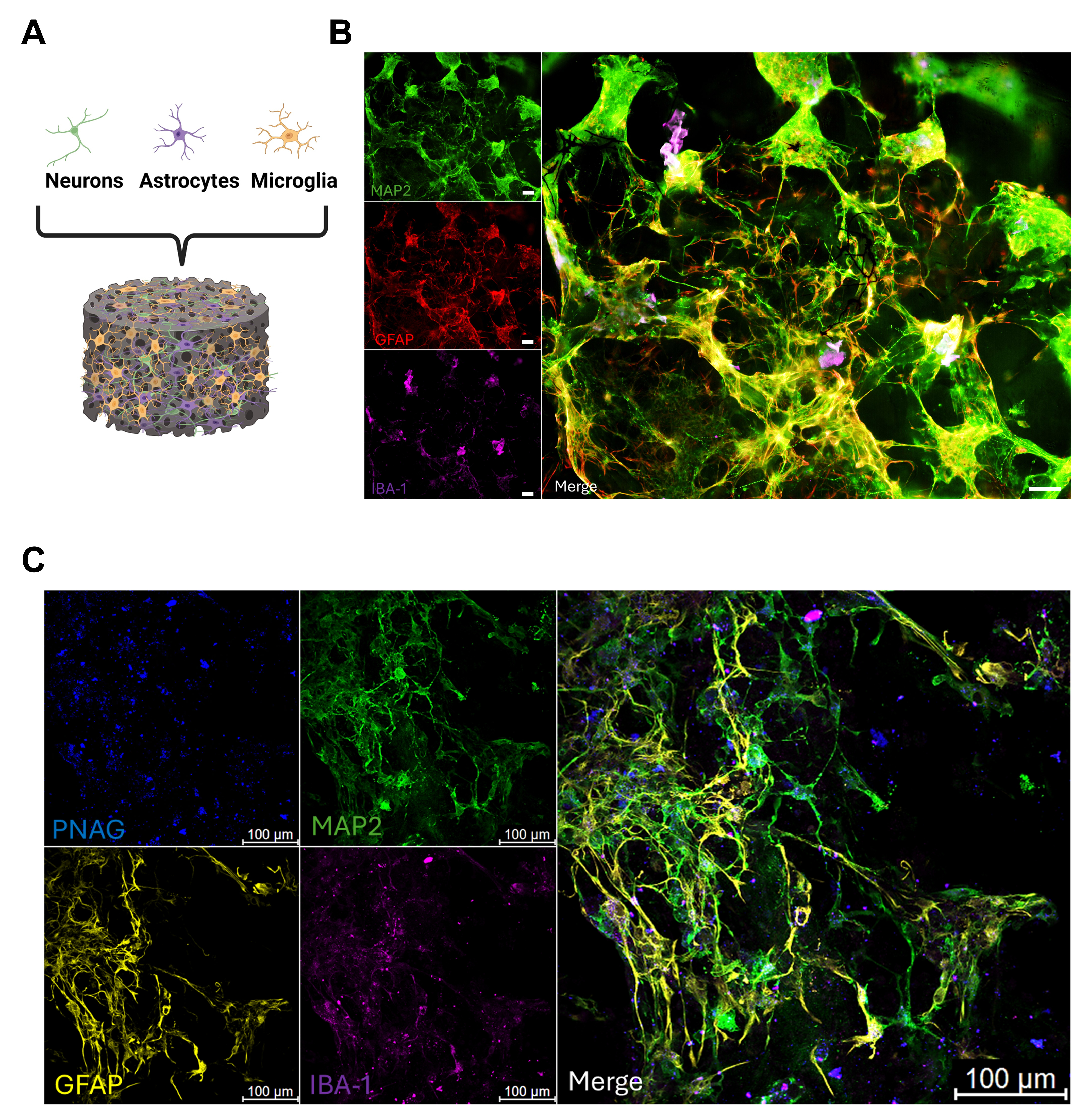

### Supplemental Figure S4

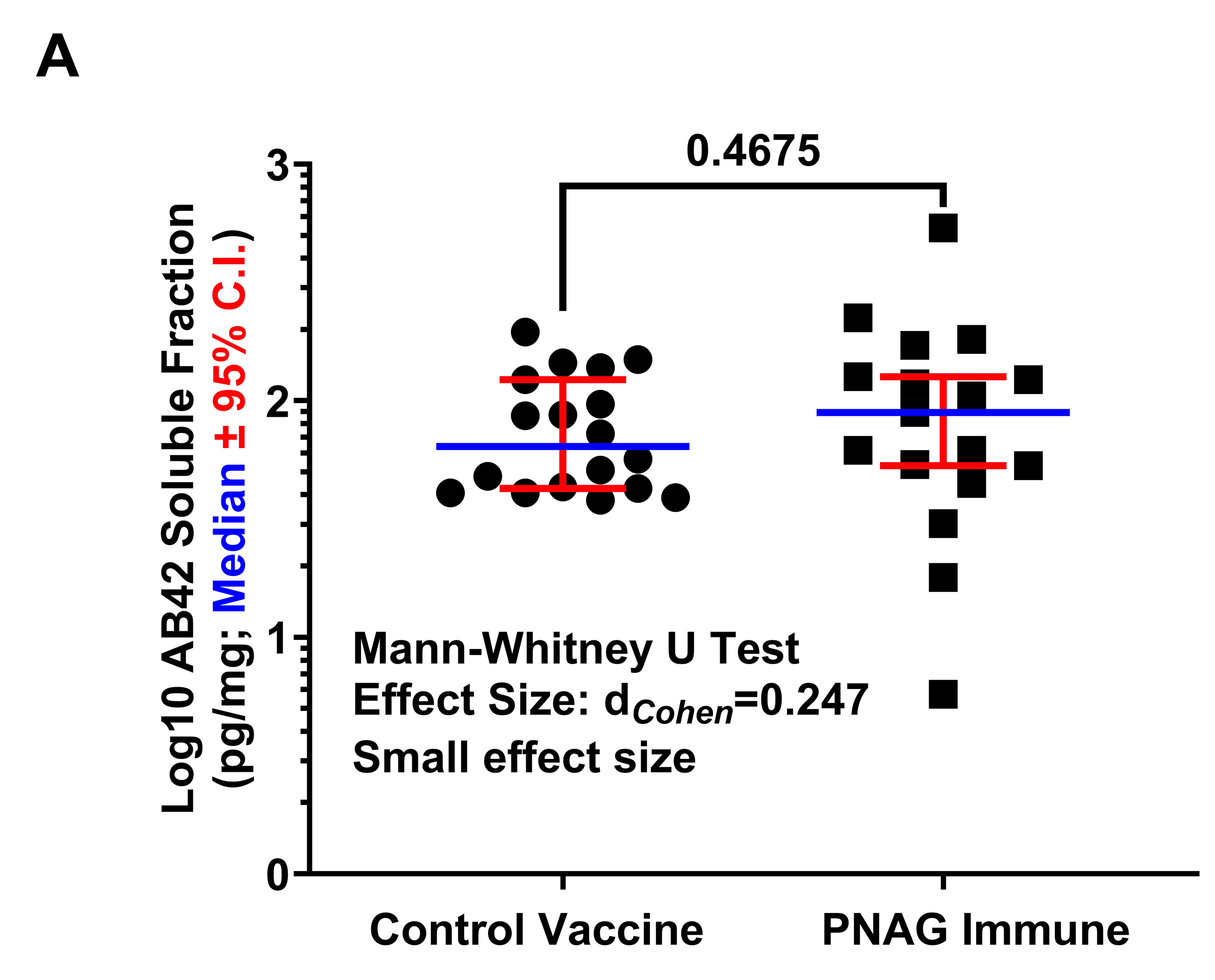

### Supplemental Figure S5

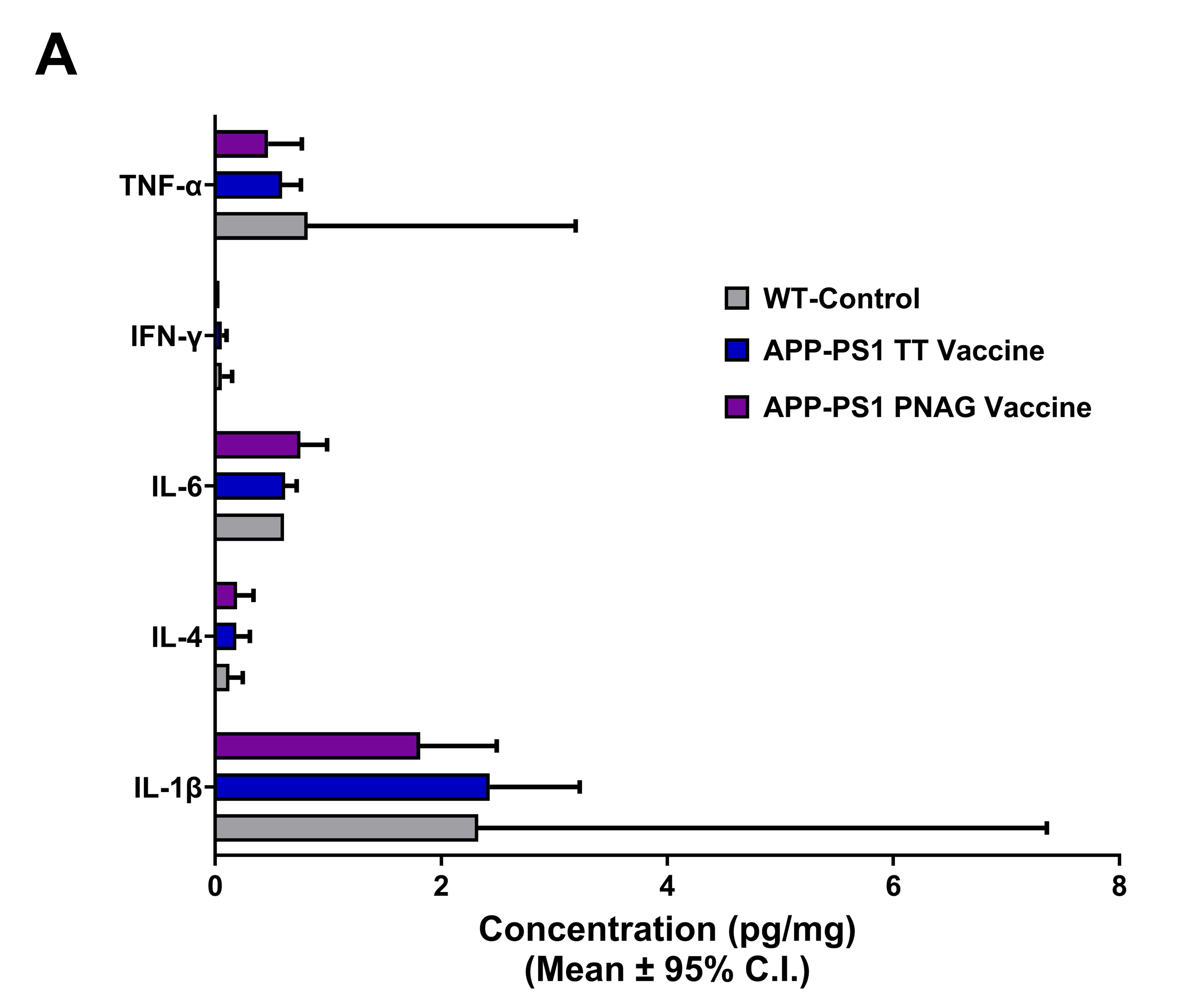

### Supplemental Figure S6

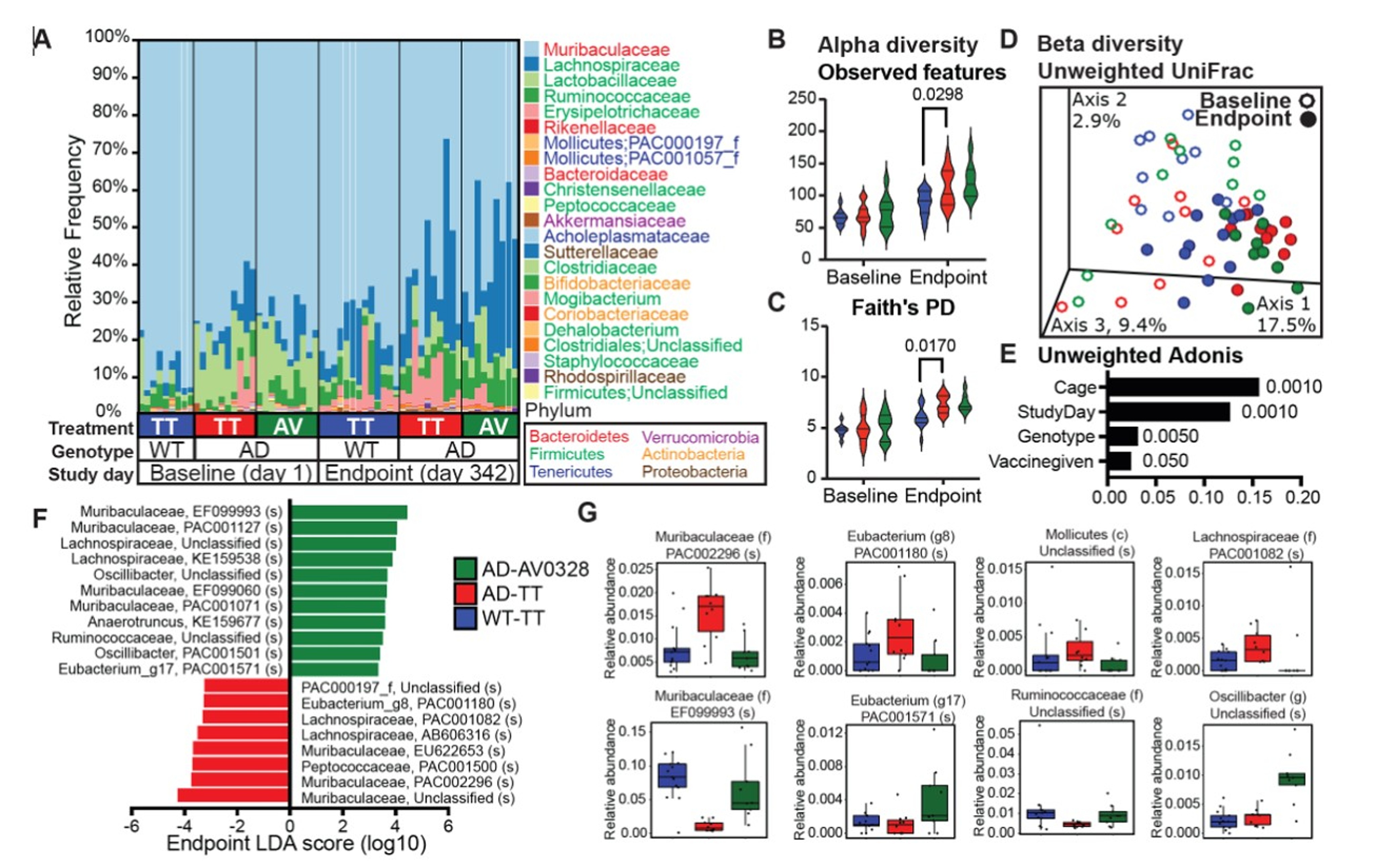
