## Supplemental Table S1 for "Persistent microbial material contributes to Alzheimer disease and is targetable by vaccination"

**Tables S1 to S2**

**Supplementary Table S1. Antibodies used for *in vitro* immunostaining.**

| Host | Antigen | Vendor | Catalog # |
|---|---|---|---|
| Mouse | CD44 | ThermoFisher | BMS150 |
| Chicken | Microtubule-associated protein 2 (MAP2) | Sigma | AB5543 |
| Mouse | Beta III tubulin (TUJ1) | Sigma | T8578 |
| Rabbit | Ionized calcium-binding adaptor molecule 1 (IBA-1) | FUJIFILM Wako | 013-27691 |
| Rat | Glial fibrillary acidic protein (GFAP) | ThermoFisher | 13-0300 |
| Rabbit | Amyloid Fibril | Abcam | ab201062 |
| Goat (Alexa 594 Conjugated) | Mouse IgG | ThermoFisher | A-11005 |
| Goat (Alexa 488 Conjugated) | Chicken IgG | ThermoFisher | A-11039 |
| Goat (Alexa 647 Conjugated) | Rabbit IgG | ThermoFisher | A-21245 |
| Goat (Alexa 405 Conjugated) | Human IgG | ThermoFisher | A48275 |
| Goat (Alexa 594 Conjugated) | Rat IgG | ThermoFisher | A-11007 |
