## Supplemental Table S2 for "Persistent microbial material contributes to Alzheimer disease and is targetable by vaccination"

**Supplementary Table S2. Primer sequences used for qRT-PCR.**

| Gene | Forward (5’→3’) | Reverse (5’→3’) |
|---|---|---|
| Interleukin 1 beta (IL-1β) | ATGATGGCTTATTACAGTGGCAA | GTCGGAGATTCGTAGCTGGA |
| Caspase-1 | TTTCCGCAAGGTTCGATTTTCA | GGCATCTGCGCTCTACCATC |
| C-X-C motif chemokine ligand 10 (CXCL10) | GTGGCATTCAAGGAGTACCTC | TGATGGCCTTCGATTCTGGATT |
| Amyloid precursor protein (APP) | CAAGCAGTGCAAGACCCATC | AGAAGGGCATCACTTACAAACTC |
| Glial fibrillary acidic protein (GFAP) | CTGGAGAGGAAGATTGAGTCGC | ACGTCAAGCTCCACATGGACCT |
| Lipocalin 2 (LCN2) | CCACCTCAGACCTGATCCCA | CCCCTGGAATTGGTTGTCCTG |
| Glyceraldehyde 3-phosphate dehydrogenase (GAPDH) | TGTGGGCATCAATGGATTTGG | ACACCATGTATTCCGGGTCAAT |

**References** (*30*)
